# Glucocorticoids desensitize hypothalamic CRH neurons to norepinephrine and somatic stress activation via rapid nitrosylation-dependent regulation of α1 adrenoreceptor trafficking

**DOI:** 10.1101/2024.07.29.605704

**Authors:** Grant L Weiss, Laura M. Harrison, Zhiying Jiang, Alyssa M. Nielsen, Maximillian S. Feygin, Sandy Nguyen, Parker S. Tirrell, Jeffrey Tasker

## Abstract

Noradrenergic afferents to hypothalamic corticotropin releasing hormone (CRH) neurons provide a major excitatory drive for somatic stress activation of the hypothalamic-pituitary-adrenal (HPA) axis. We showed that glucocorticoids rapidly desensitize CRH neurons to norepinephrine and suppress inflammation-induced HPA activation via a glucocorticoid receptor- and endocytosis-dependent mechanism. Here, we show that α1 adrenoreceptor (ARα1) trafficking is regulated by convergent glucocorticoid and nitric oxide synthase signaling mechanisms. Live-cell imaging of ARα1b-eGFP-expressing hypothalamic cells revealed rapid corticosterone-stimulated redistribution of internalized ARα1 from rapid recycling endosomes to late endosomes and lysosomes via a nitrosylation-regulated mechanism. Proximity assay demonstrated interaction of glucocorticoid receptors with ARα1b and β-arrestin, and showed corticosterone blockade of norepinephrine-stimulated ARα1b/β-arrestin interaction, which may prevent ARα1b from entering the rapid recycling endosomal pathway. These findings demonstrate a rapid glucocorticoid regulation of G protein-coupled receptor trafficking and provide a molecular mechanism for rapid glucocorticoid desensitization of noradrenergic signaling in CRH neurons.

## INTRODUCTION

Increased circulating glucocorticoids induced by activation of the hypothalamic-pituitary-adrenal (HPA) axis play an important role in the metabolic, immunoregulatory, and cognitive effects of stress ^1,2^. Rapid glucocorticoid-mediated negative feedback regulation of the corticotropin releasing hormone (CRH) neurons in the hypothalamic paraventricular nucleus (PVN) contributes to the termination of the HPA response ^3^. While the mechanism responsible for negative feedback regulation of the HPA axis has been classically attributed to transcriptional regulation by glucocorticoid receptors ^4,5^, several studies have shown that glucocorticoids rapidly inhibit PVN CRH neurons (Di et al., 2003; Tasker et al., 2006) and pituitary corticotrophs ^9^ to mediate fast negative feedback (Evanson et al., 2010). Noradrenergic afferents to the CRH neurons provide a primary excitatory drive for the somatic stress activation of the HPA axis ^11^ via activation of α1 adrenergic receptors (ARα1) ^12–15^. We recently reported that glucocorticoid rapidly desensitizes PVN CRH neurons to ARα1 activation by norepinephrine (NE), which suppresses the HPA response to somatic stress, but not psychological stress, due to the NE dependence of somatic stress activation of the HPA axis ^11,16^. A main source of noradrenergic input to PVN CRH neurons is the brainstem nucleus of the solitary tract (NTS) ^17^. The NTS is activated by inputs from the vagal nerve and relays to the forebrain somatic interoceptive stress signals such as organ damage, metabolic challenge, and immune activation ^18^. Stimulation of an immune response by systemic lipopolysaccharide (LPS), for example, activates the HPA axis by way of the NTS noradrenergic system ^19,20^. Our recent findings suggest that prior priming by acute stress exposure attenuates subsequent LPS activation of the HPA axis via a rapid glucocorticoid-induced desensitization of the CRH neurons to NE activation of ARα1 ^21^, but the cellular and molecular mechanisms of this desensitization are not known.

The stress desensitization of PVN CRH neurons to NE is dependent on the ligand-induced internalization of ARα1 ^21^. ARα1 is internalized by clathrin-mediated endocytosis and is trafficked back to the membrane via recycling endosomes, a process that involves multiple trafficking proteins and that establishes a steady state of surface receptors and NE sensitivity ^22^. Initial steps of endocytosis following ligand binding and G protein activation include receptor phosphorylation, recruitment of β-arrestin to the phosphorylated receptor, and engagement of clathrin and dynamin to execute endocytosis. β-arrestin and the internalized ARα1 undergo post-translational modifications, including nitrosylation and ubiquitination, which regulate trafficking through the cell’s endosomal pathways ^23,24^. Here, we tested the cellular mechanism by which glucocorticoids desensitize NE signaling. We found that corticosterone (CORT) desensitizes CRH neurons to NE in a process involving nitric oxide synthesis (NOS)-dependent protein S-nitrosylation that rapidly depletes the cells of surface adrenoreceptors by re-routing the ARα1 out of the rapid membrane recycling endosomal pathway and into late endosomes and lysosomes.

## RESULTS

### Corticosterone causes a rapid, long-lasting desensitization of ARα1

Our previous study (Chen et al., 2019) showed that in *ex vivo* brain slices, PVN CRH neurons in the male mouse respond to NE with an increase in excitatory postsynaptic currents (EPSCs) that is mediated by postsynaptic ARα1 activation and stimulation of retrograde glial-neuronal circuits by dendritic volume transmission (see diagram in Fig. 1A). A follow-up study showed that the PVN CRH neuron excitatory response to NE is suppressed by 5-10 min of preincubation of brain slices from unstressed mice in CORT (2 μM) (Fig. 1B) and by prior restraint stress-induced endogenous glucocorticoid exposure *in vivo* (Fig. 1C) ^21^. The rapid glucocorticoid-induced desensitization of CRH neurons to NE was blocked by a dynamin inhibitor, indicating that it is dependent on endocytosis (Fig. 1C).

**Figure 1.**
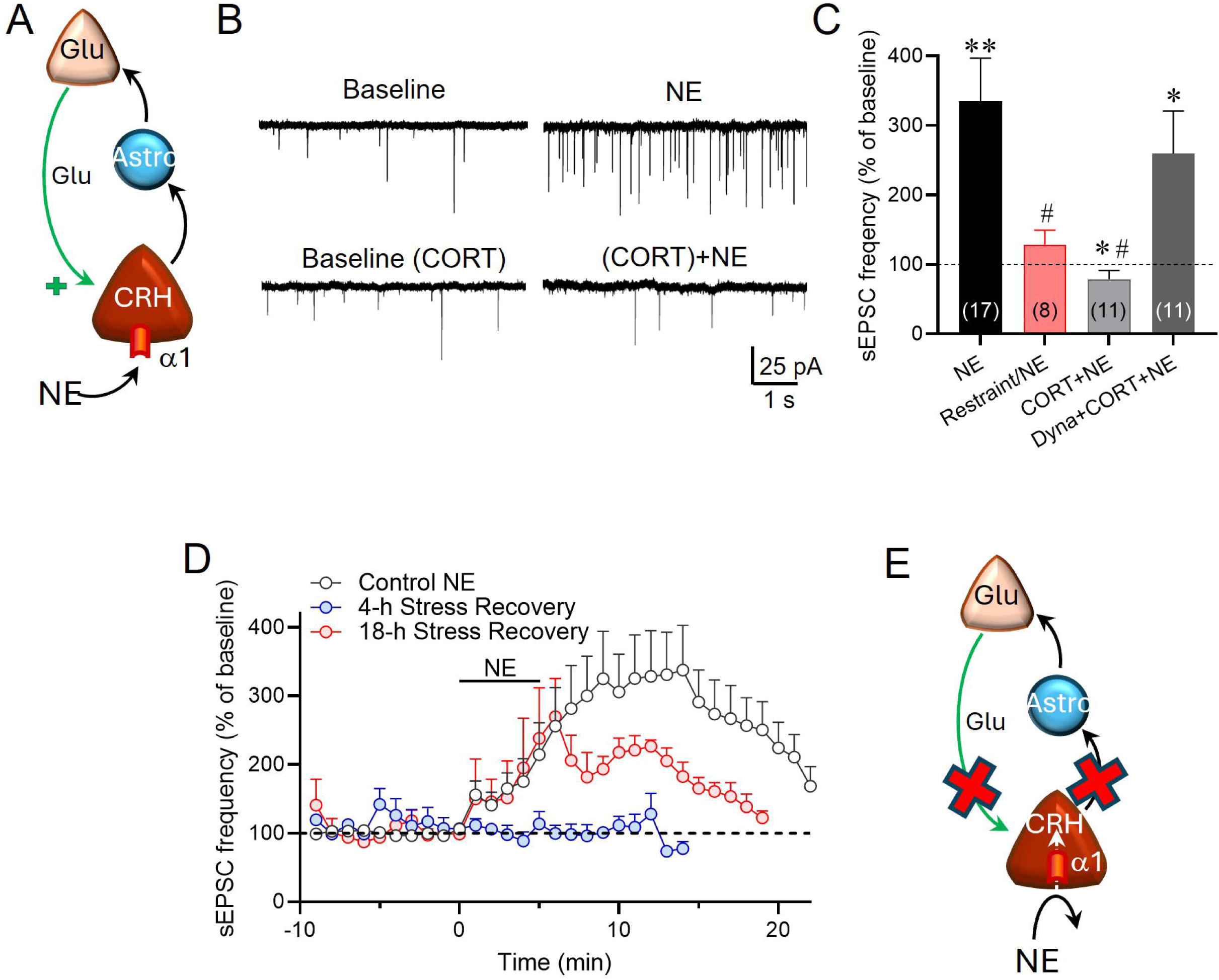
Long-lasting glucocorticoid desensitization of NE ARα1-induced increase in sEPSC frequency in CRH neurons via a dynamin-dependent mechanism. **A.** Model of retrograde neuronal-glial signaling response triggered in CRH neurons by NE, based on Chen et al., 2019. **B.** Representative traces from our previous study of the sEPSC response in PVN CRH neurons to NE (100 µM) without pretreatment (Baseline/NE) (upper trace) and in the presence of 1 μM corticosterone (Baseline (CORT)/NE (CORT), 10 min) (lower trace). **C.** Summary graph of the stress (Restraint) and CORT (CORT) suppression of the NE-induced increase in sEPSC frequency and the rescue of the NE response by blocking endocytosis with the dynamin inhibitor dynasore (80 μM, 10 min) (Dyna+CORT+NE). **D.** Time histogram of the NE-induced increase in sEPSC frequency in CRH neurons over time in slices after 4 h or 18 h recovery from a 30-min acute restraint stress (4-h and 8-h Stress Recovery), compared to the NE response following no prior stress (Control NE, taken from ^21^). Complete desensitization of the NE response was seen after a recovery period of 4 h following restraint and the response was partially restored after 18 h of recovery. Panels B and C and the Control NE graph in D were modified from ^21^ with permission.

Here, we tested the duration of the stress-induced desensitization of CRH neurons to NE by allowing the mice to recover from a 30-min restraint stress in their home cage for 4 h and 18 h prior to sacrifice and brain slice preparation. In animals allowed to recover from acute restraint for 4 h, the PVN CRH neurons remained insensitive to NE (100 µM), whereas the EPSC response to NE was partially restored 18 h after the restraint stress (Fig. 1D). This indicated that the stress-induced desensitization of PVN CRH neurons to activation by NE takes longer than 18 h to fully reverse and suggests a surprisingly long-lasting diminished CRH neuron sensitivity to NE and to adrenoreceptor activation of the HPA axis following exposure to acute stress ^21^. Thus, acute glucocorticoid exposure causes a long-lasting desensitization of CRH neurons to NE via a mechanism with rapid onset, which is dependent on dynamin-dependent endocytosis of the ARα1 (Fig. 1E).

ARα1b has been reported to be the dominant α1 adrenoreceptor subtype in CRH neurons ^25^, and we previously reported evidence of ARα1b internalization in fixed N42 cell cultures ^21^. Here, we used live-cell imaging to follow the internalization and trafficking of the ARα1b through the endosomal pathway. We used the N42 immortalized hypothalamic cell line ^26^, which we transiently transfected with an pEGFP-ARα1b construct and imaged live by confocal microscopy to visualize ARα1b receptor trafficking. A low NE concentration (1 μM) caused ARα1b internalization, which was seen by a decrease in membrane pEGFP-ARα1b and an increase in cytosolic pEGFP-ARα1b fluorescence, which reached a peak at ∼30 min (Fig. 2A and B), a time course that coincided approximately with that of the NE-induced increase in sEPSC frequency (see Fig. 1). Following 30 min of NE application, CORT (2 μM) was co-applied with NE, which induced a rapid increase in the ARα1b internalization over the NE-induced steady-state internalization. The onset of the CORT effect occurred within minutes and peaked within 10 min of its introduction into the chamber (Fig. 2A and B), which was consistent with a rapid CORT-induced desensitization of CRH neurons to NE. The combination of NE and CORT caused significantly more internalization than NE alone (Fig. 2B, C). The bulk of the NE- and CORT-induced increases in cytosolic ARα1b appeared to be directed mainly to a single “hot-spot” in the cells (arrows in Fig. 2A), which we targeted for quantitation and used to map the trafficking of the ARα1b receptor in the endocytic pathway with endosomal markers. The dynamic change in ARa1b distribution within the cell during NE and NE+CORT application can be followed in real time in the supplemental video 1.

**Figure 2.**
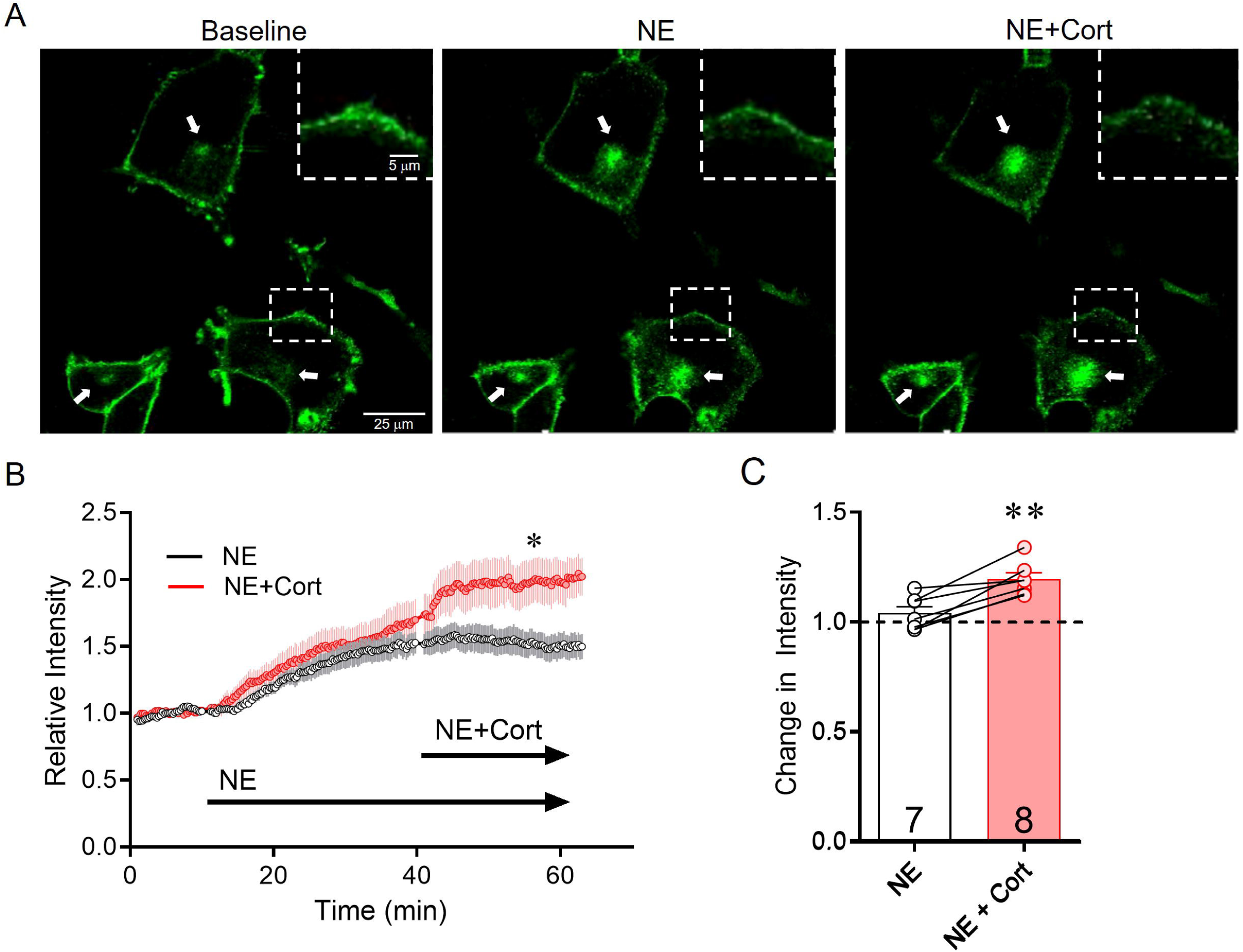
Cort causes a rapid increase in cytosolic ARα1b accumulation in a distinct subcellular “hot spot”. **A.** In N42 cells transiently transfected with ARα1b-eGFP, ARα1b is mostly localized to the membrane at baseline, but redistributes from the membrane to internal ‘hotspots’ (arrows) after 30 min of NE application (NE) and further after 20 min of application of NE and CORT (NE+Cort). Membrane region in the dashed box is enlarged in the upper insets. **B**. Time course of increase in intracellular ARα1b-eGFP. Following plateau of the increase in the fluorescence signal by NE (Vehicle), the co-application of CORT caused a rapid, additional increase in cytosolic ARα1b that plateaued within 10 min (*, p<0.05, t-test vs NE). **C**. Relative increase in cytosolic intensity of ARα1b-eGFP compared to NE alone (NE) following the addition of CORT (NE+Cort). (**, p<0.01, paired t-test vs. NE).

### Cort redirects AR**α**1b trafficking from rapid recycling endosomes to late endosomes

Receptor-mediated endocytosis forms early endosomes, which either cycle to the plasma membrane as recycling endosomes to replenish surface receptors or develop into late endosomes and are routed out of the recycling pathways. Specific Rab GTPases regulate distinct stages of endosomal trafficking ^27^. By co-transfecting the N42 cells with pEGFP-ARα1b and constructs for different dsRed-tagged Rab proteins, we were able to use Förster resonance energy transfer (FRET) analysis to detect with 10-nm resolution the colocalization of the ARα1b with the specific markers of different endosomal compartments. We tested for ARα1b association with the early endosomal marker Rab5, the late endosomal marker Rab 7, the rapid recycling endosomal marker Rab4a, and the slow recycling endosomal marker Rab 11 ^28^.

Cells treated with NE (1 µM) were first analyzed for an increase in ARα1b FRET with dsRed-tagged Rab5, a marker of newly internalized cargo being transported to the early endosome. Norepinephrine caused the expected increase in ARα1b localization in the early endosome, serving as a control for ligand-mediated endocytosis of the adrenoreceptor. However, subsequent CORT application failed to increase colocalization of ARα1b with Rab5 compared to that caused by NE alone (Fig. 3A). This indicated that the CORT-induced increase in ARα1b intracellular concentration was mediated by an accumulation of ARα1b in the cytosol following standard ligand-dependent internalization and not by a CORT-induced increase in ARα1b endocytosis. We next tested whether CORT caused an increase in the receptor trafficking to the late endosome using ARα1b FRET with the late endosomal marker Rab7. Norepinephrine alone caused an increase in the ARα1b-Rab7 FRET signal, and subsequent CORT treatment increased the FRET signal compared to that seen with NE alone (Fig. 3B). This indicated that CORT caused an increase in the NE-induced ARα1b trafficking to the late endosome and suggests that the CORT-induced desensitization is not associated with an increase in the amount of ARα1b undergoing internalization, but rather that it causes an increase in the receptor trafficking into the late endosomal pathway. A possible explanation for the accumulation of ARα1b in the late endosome and loss of sensitivity to NE could be a decrease in the trafficking of the receptor back to the plasma membrane.

**Figure 3.**
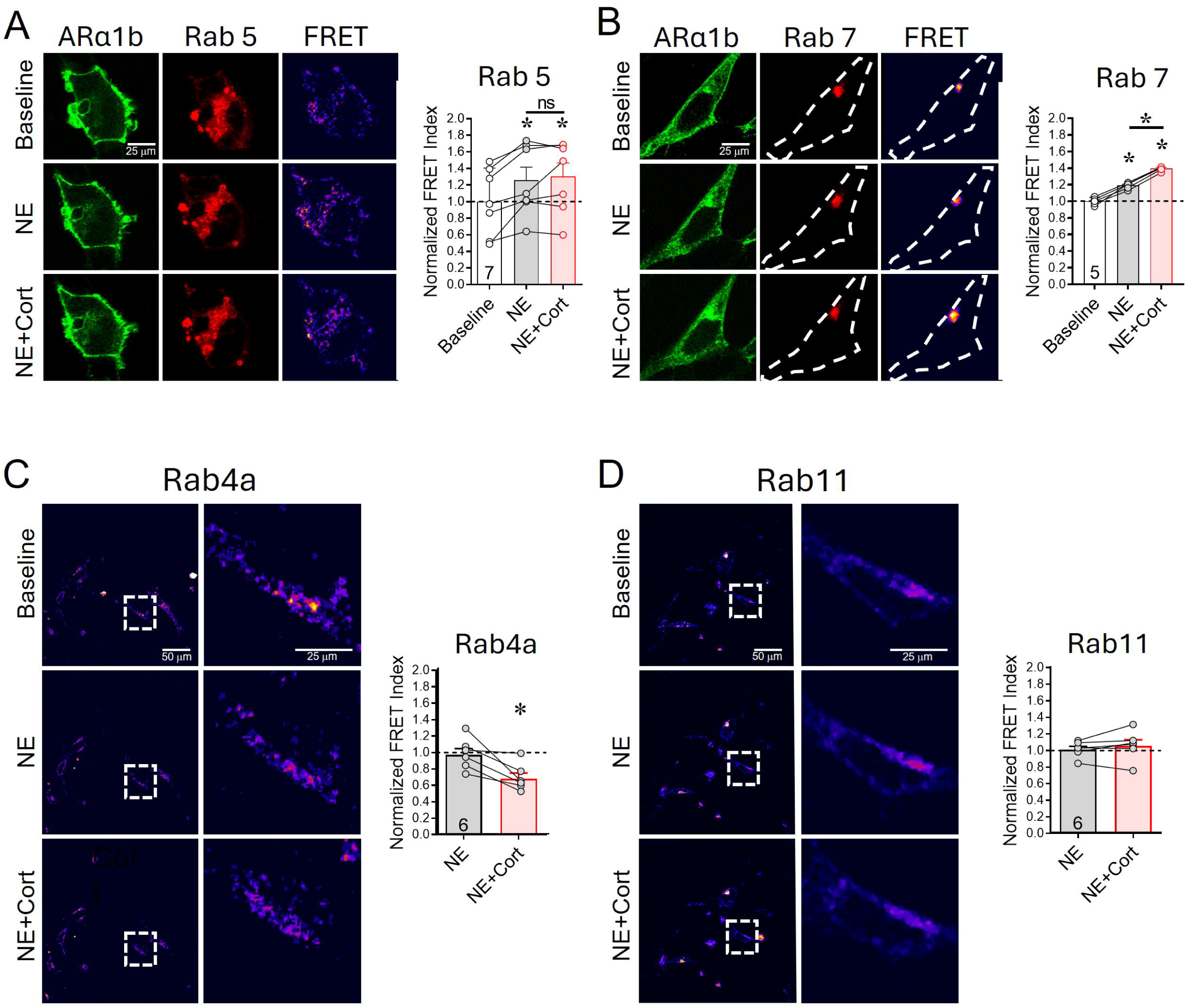
Cort redirects ARα1b trafficking from the rapid recycling endosome to the late endosome. **A.** N42 cells expressing ARα1b-eGFP (ARα1b) and Rab5-dsRed (Rab 5) and images of calculated FRET ratios (FRET) after treatments with NE and NE+Cort. **Right**: Mean (+/- SEM) of FRET signal in cells treated with NE (1 µM) and NE+Cort showing increased FRET between ARα1b and Rab5 induced by NE, but no further increase after subsequent Cort application (2 µM) (repeated measures ANOVA, Tukey’s post-hoc, * p < 0.05 vs baseline; n.s., p=0.7423). **B**. N42 cells expressing ARα1b-eGFP (ARα1b) and Rab7-dsRed (Rab 7) and images of calculated FRET ratios (FRET) after treatments with NE and NE+Cort. **Right**: Mean (+/- SEM) of FRET signal in cells treated with NE (1 µM) and NE+Cort, showing increased FRET between ARα1b and Rab7 with NE, and a further increase in FRET ratio with subsequent Cort application (2 µM) (repeated measures ANOVA, Tukey’s post-hoc, * p<0.05). **C.** Images of calculated FRET ratios of N42 cells expressing ARα1b-GFP and Rab4a-dsRed after NE and Cort treatments. Images in the right column show the areas in the squares at higher magnification. **Right:** Mean (+/- SEM) of FRET signal showing no significant change in ARα1b/Rab4a interaction after treatment with NE (1 µM) and a decrease in FRET ratio with subsequent Cort application (2 µM) (repeated measures ANOVA, * p<0.05 vs NE). **D.** Images of calculated FRET ratios of N42 cells expressing ARα1b-GFP and Rab11-dsRed after NE and NE+Cort treatments. Images in the right column show the areas in the squares at higher magnification **Right:** Mean (+/- SEM) of FRET signal in NE and NE+Cort, normalized to baseline. There were no significant changes to the ARα1b-Rab 11 FRET ratio after application of either NE (1 µM) or NE+Cort (2 µM) (repeated measures ANOVA, p=0.2895).

We tested this using FRET analysis with the rapid recycling endosomal marker Rab4a and with the slow recycling endosomal marker Rab11. Norepinephrine did not cause a change in the ARα1b-Rab4a FRET signal, and the addition of CORT decreased the ARα1b-Rab4a signal compared to NE alone (Fig. 3C), indicating reduced ARa1b association with Rab4a and suggesting a reduction in trafficking of the ARα1b through the rapid recycling endosome. Finally, we tested for changes in the ARα1b interaction with the slow recycling endosomal marker Rab11. Neither NE nor NE+CORT had any effect on the ARα1b-Rab 11 FRET signal (Fig. 3D), suggesting that neither treatment increases the slow recycling of ARα1b to the membrane within the timeframe of the desensitization to NE. Together, these data indicate that CORT influences ARα1b trafficking by redirecting the receptor away from rapidly recycling endosomes and into late endosomes. This would be expected to deplete the membrane of adrenoreceptors and provides, therefore, a molecular substrate for the glucocorticoid-induced desensitization to NE observed in *ex vivo* recordings.

### Cort causes AR**α**1b trafficking to the lysosome

To determine whether CORT causes ARα1b to travel further down the endocytic pathway from late endosomes to lysosomes, we tested for ARα1b trafficking to the lysosome by imaging the co-localization of pEGFP-ARα1b with the lysosomal marker LAMP1 conjugated to red fluorescent protein (LAMP1-RFP). Application of NE (1 μM) caused an increase in ARα1b-LAMP1 co-staining (Fig. 4A-C), indicating an increase in the co-localization of the two proteins. After 30 min of NE application, CORT co-application (2 µM) further increased the ARα1b-LAMP1 co-staining (Fig. 4A-C), which revealed a CORT facilitation of ARα1b trafficking to the lysosome for degradation. However, a large proportion of the ARα1b was not co-stained with the LAMP1 marker, including the CORT-induced ARα1b “hot spot”, resulting in a low Pearson’s correlation of ARα1b and LAMP1 localization (maximum Pearson’s coefficient ∼0.5) (Fig. 4B), which suggests that, on this time scale (∼50 min), most of the ARα1b receptor was not trafficked to the lysosome for degradation, but rather was sequestered within the cell.

**Figure 4.**
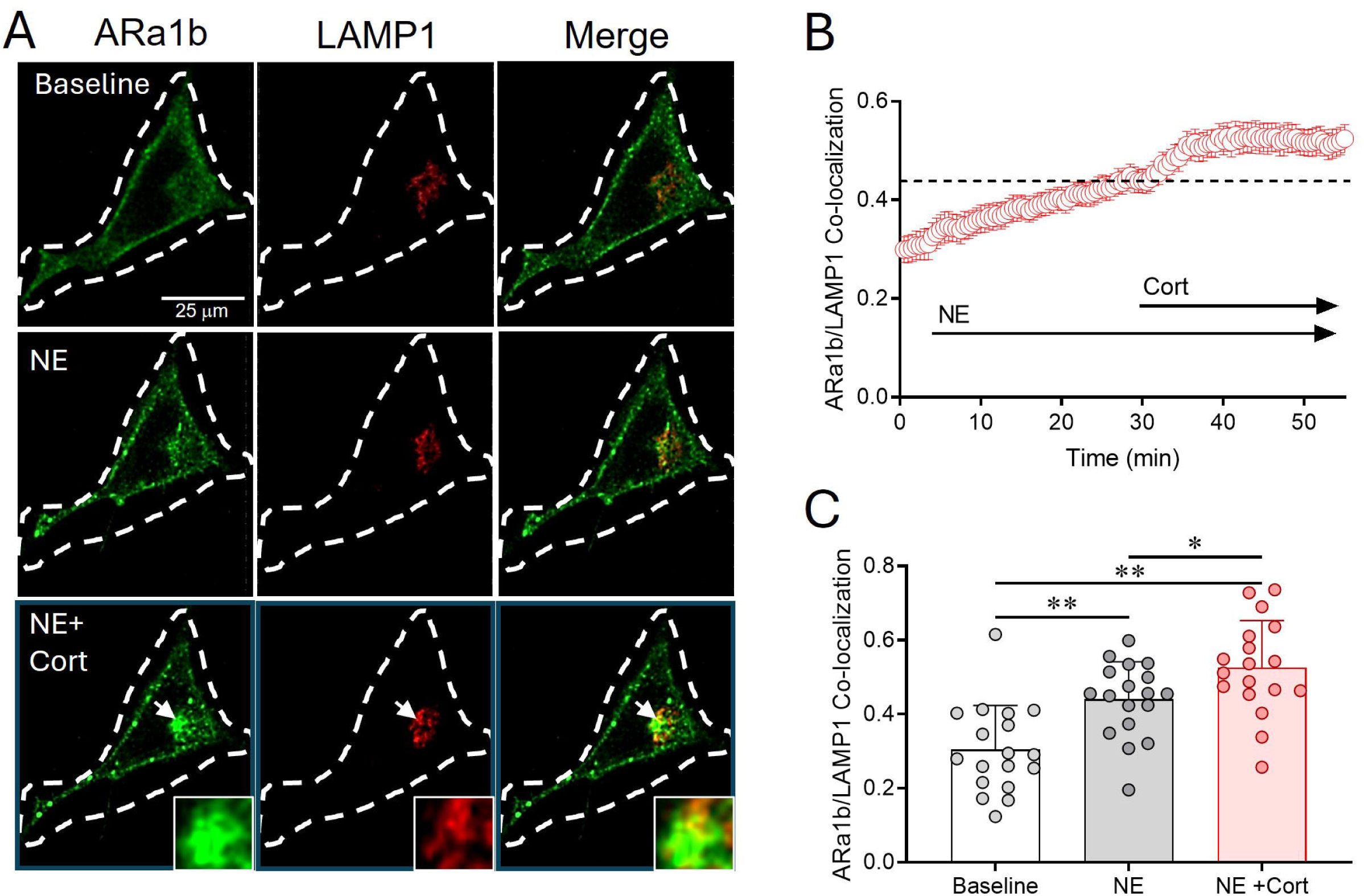
ARα1b trafficking to lysosomes. **A.** ARα1b-eGFP and LAMP1-RFP were co-transfected into N42 cells, and the cells were treated with NE for 30 min, followed by NE + CORT for 20 min. Arrow: CORT-facilitated internalized ARα1b co-localized with LAMP1, a lysosomal marker. **B.** Time lapse quantification of co-localization by Pearson’s coefficient. **C.** Co-localization averages of baseline and the last 3 minutes of NE and NE +CORT treatment (Mean +/- SEM, ANOVA, Tukey’s post-hoc, * p<0.05, ** p<0.01).

### Blocking AR**α**1 reverses Cort-induced AR**α**1b trafficking

We found previously that CORT alone without prior NE application has no effect on ARα1b internalization in N42 cells and that inhibiting ARα1 with prazosin during CORT application in hypothalamic slices blocked the glucocorticoid-induced desensitization of CRH neurons to NE ^21^, suggesting that glucocorticoid-induced desensitization to NE requires ligand (NE)-induced ARα1 internalization. Here, we tested whether N42 cells in which CORT had already caused increased ARα1b cytosolic accumulation would recover after blocking ARα1 with prazosin. As described above (see Fig. 1 and Fig. 2A-C), 1 μM NE caused ARα1b accumulation in the cytosol that was further increased by CORT (2 μM). The addition of the ARα1 antagonist prazosin (1 μM) caused the internalized ARα1b to return to near baseline levels within 10 min (Fig. 5A, B). These data suggest that blocking ARα1 reverses glucocorticoid-induced desensitization to NE.

**Figure 5.**
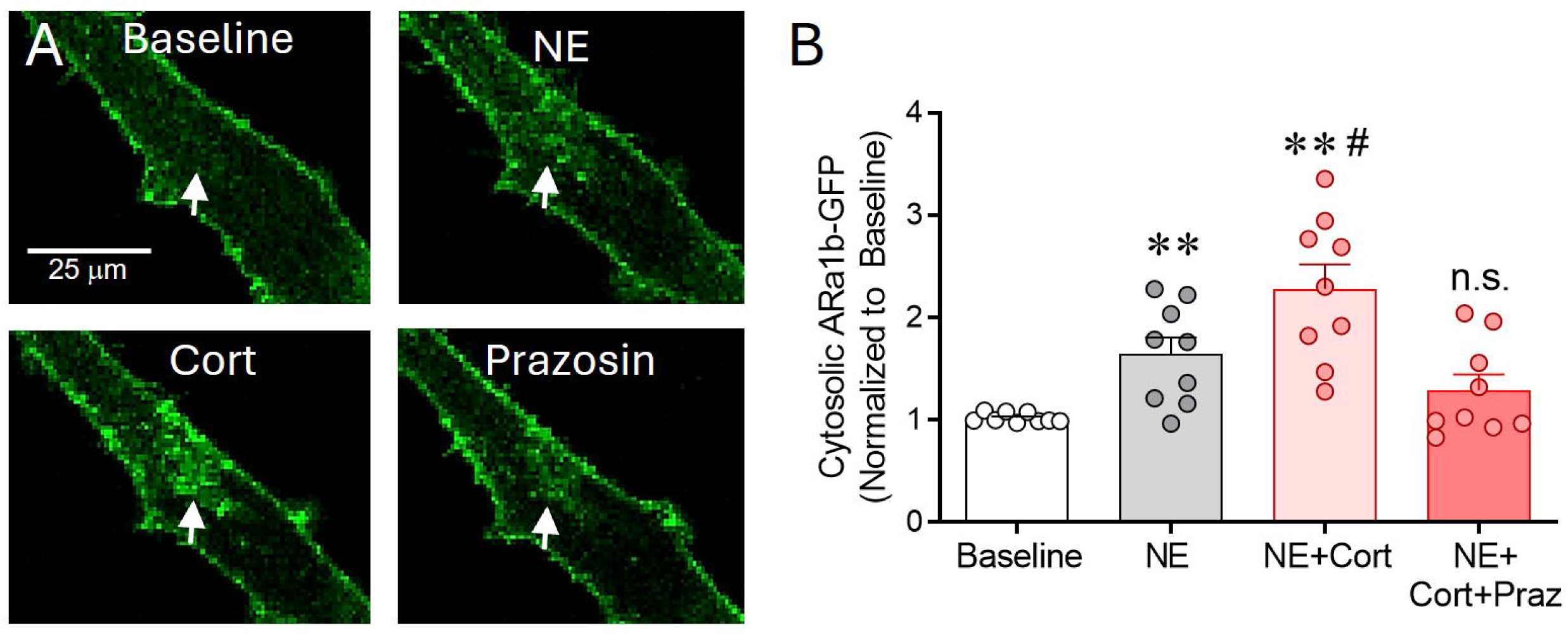
Reversal of NE- and Cort-induced ARα1b trafficking. **A**) ARα1b-EGFP-transfected N42 cells were treated with NE for 30 min, followed by the addition of Cort for 20 min and finally the addition of prazosin for 30 min. Arrows: ARα1b-EGFP hot spot. **B**) Normalized cytosolic fluorescence (ANOVA, Tukey’s post-hoc, * p<0.05 vs. baseline, # p<0.05 vs. NE). The ARα1b hot spot returned to near baseline after 30 min prazosin co-application (n.s., p=0.35 vs baseline).

### Nitrosylation dependence of the CRH neuron response to norepinephrine

Since GPCR internalization and trafficking are dependent on S-nitrosylation of proteins associated with receptor trafficking, including β-arrestin ^24^, we next tested whether the CORT control of ARα1 trafficking is regulated by nitric oxide synthase (NOS) activity. We first tested this with whole-cell recordings of the NE response in CRH neurons in *ex vivo* hypothalamic slices. Blocking NOS activity with the broad-spectrum NOS inhibitor N(ω)-nitro-L-arginine methyl ester (L-NAME, 50 μM) inhibited the NE-induced increase in sEPSC frequency (Fig. 6A, B), which was similar to the glucocorticoid inhibition of the NE-induced increase in sEPSC frequency in CRH neurons (see Fig. 1). Since NOS induces both the production of the gaseous messenger NO and post-translational S-nitrosylation of target proteins, we tested for these two NOS-dependent mechanisms pharmacologically. We first blocked the NO receptor, soluble guanylyl cyclase, with 1H-[1,2,4]oxadiazolo[4,3,-a]quinoxalin-1-one (ODQ) (100 μM), but this failed to inhibit the NE-induced increase in sEPSC frequency in the CRH neurons (Fig. 6A, B). This confirmed our previous finding that NE does not induce an NO retrograde signal at glutamate synapses on the CRH neurons ^15^. We next tested the effect of the S-nitrosylation inhibitor N-ethylmaleimide (NEM) (50 µM) on the NE response. NEM completely blocked the NE-induced increase in sEPSC frequency in the CRH neurons (Fig. 6A, B). These findings suggested that the NOS dependence of the NE activation of excitatory inputs to the CRH neurons is due to S-nitrosylation of target proteins in the CRH neurons, and not to the release of NO as a transmitter.

**Figure 6.**
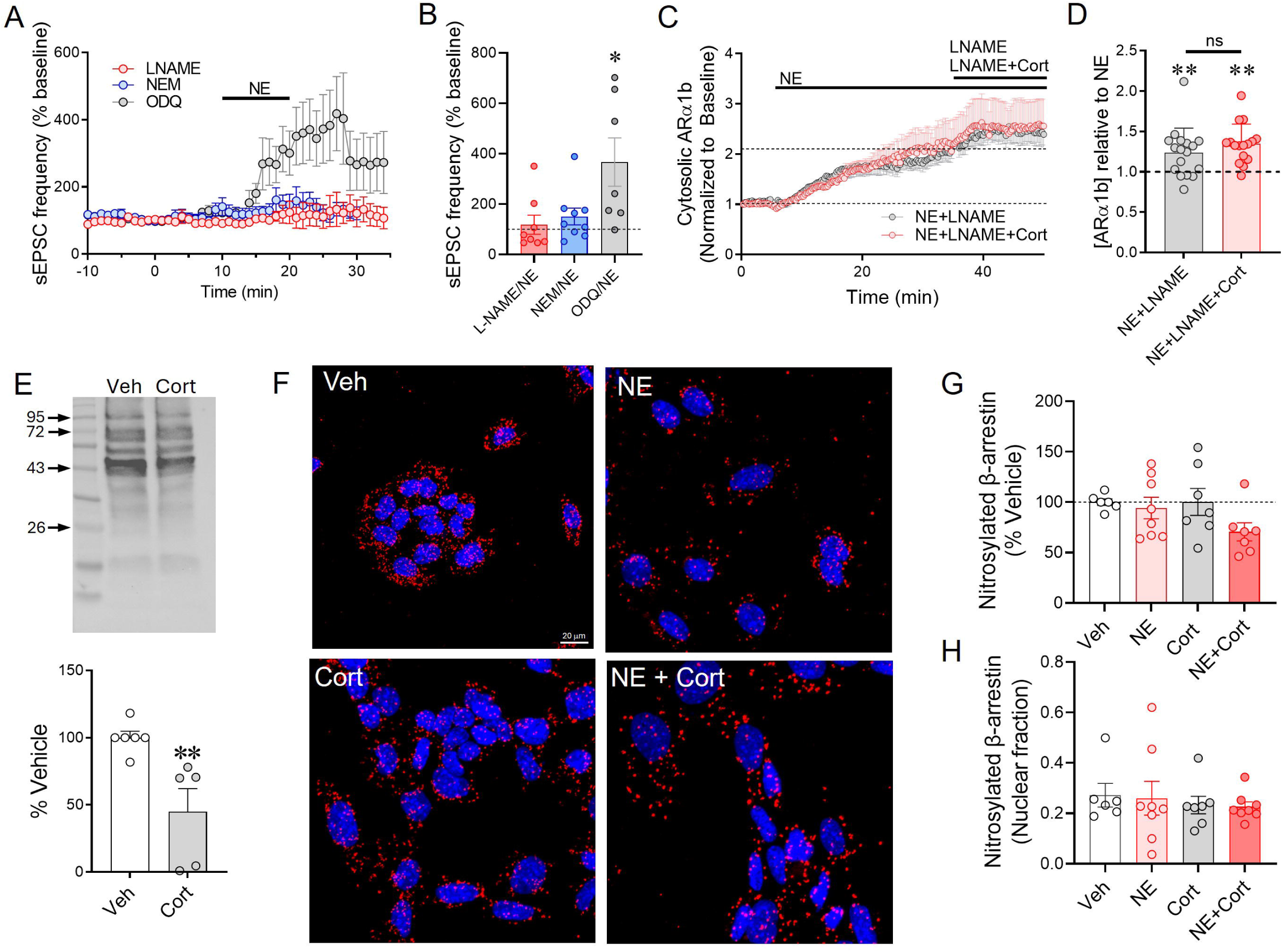
CORT regulation of protein S-nitrosylation. **A.** Nitrosylation dependence of the NE-induced increase in sEPSCs in PVN CRH neurons. The NE-induced increase in sEPSC frequency was blocked by the nitric oxide synthase (NOS) inhibitor L-NAME and the S-nitrosylation inhibitor NEM, but not by the soluble guanylyl cyclase inhibitor ODQ. **B.** Summary graph of the effects of L-NAME, NEM, and ODQ on the NE-induced increase in sEPSC frequency. **C.** Changes over time in the cytosolic intensity of ARα1b-eGFP in N42 cells after treatment with NE and L-NAME (NE+LNAME) and NE, L-NAME and CORT (NE+LNAME+Cort). **D.** Summary graph of the changes in ARα1b-EGFP fluorescence intensity indicative of internalization in NE+L-NAME and NE+L-NAME+Cort, normalized to the baseline fluorescence intensity (** p<0.01 compared to baseline). Cort and L-NAME co-application was not significantly different from L-NAME alone (p=0.2009, Students *t*-test). Cytosolic ARα1b measurements in L-NAME and L-NAME+Cort were taken at 40-45 min and compared to measurements in NE alone at 30-35 min. **E.** Western blot analysis of whole-cell lysate (cytosolic fraction) from N42 cells treated with vehicle or 2 µM Cort followed by TMT switch assay for total nitrosylated protein. Cort-treated cells showed a significant decrease in total nitrosylated protein [t(9) = 3.368, p = 0.0083, Student’s *t*-test, n=5-6]. **F.** Images of β-arrestin1/2 nitrosylation in N42 cells treated with NE, Cort, or NE+Cort for 20 min. The biotin-switch assay was performed, followed by proximity ligation assay (PLA) with antibodies to biotin and β-arrestin1/2. Maximum intensity projections of z-stacks. Red dots = PLA signal representing nitrosylated β-arrestin1/2; blue = DAPI. **G.** Nitrosylation of β-arrestin1/2 by NE, Cort, and NE+Cort relative to vehicle. There were no significant drug effects [One-Way ANOVA, F (3, 24) = 1.861; p = 0.1632 n = 6-8 fields from 4 separate assays]. **H.** None of the drug treatments altered the nuclear localization of nitrosylated β-arrestin 1/2 [One-Way ANOVA, F (3, 25) = 0.2101; p = 0.8885, n = 6-8 fields from 4 separate assays].

Several proteins involved in GPCR trafficking are regulated by nitrosylation, including G protein-coupled receptor kinase 2 (GRK2), β-arrestin, and clathrin ^24,29^. We first tested for the nitrosylation dependence of intracellular trafficking of the ARα1b with live-cell imaging in N42 cells. We found that N42 cells express endothelial NOS and inducible NOS, but not neuronal NOS (Supplemental Fig. 1). Co-application of the NOS inhibitor L-NAME (50 μM) with NE (1 μM) caused an increase in the ARα1b internalization over that induced by NE application alone (Fig. 6C, D), suggesting that inhibiting S-nitrosylation enhanced ligand-dependent internalization/trafficking of the receptor. Co-application of CORT (2 µM) with L-NAME failed to increase the ARα1b cytosolic concentration compared to L-NAME alone (Fig. 6C, D), indicating that CORT did not cause any additional intracellular ARα1b accumulation. This suggested, therefore, that blocking S-nitrosylation caused an occlusion of the CORT effect, revealing a convergence of the two signaling mechanisms and opposite effects of CORT (i.e., increased internal ARα1b accumulation) and S-nitrosylation (i.e., decreased internal ARα1b accumulation).

We next tested for changes in S-nitrosylation in the N42 cells with a biochemical approach using Western blot analysis and a proximity ligation assay following the application of a modified biotin-switch procedure. The biotin-switch procedure consists of cleaving the NO groups on cysteine residues of nitrosylated proteins and labeling these thiols with biotin or another tag such as a tandem mass tag (TMT) ^30^. Western blot of TMT-tagged proteins in the cytosolic fraction of N42 cells showed a significant decrease in total S-nitrosylation with a 20-min application of CORT (2 µM) (Fig. 6E). This was likely due to activation of the glucocorticoid receptor because we found a similar decrease in total S-nitrosylation in N42 cells with the synthetic glucocorticoid dexamethasone (1 µM) (data not shown).

Western blot analysis revealed a strong nitrosylation signal at a molecular weight of about 50 kD, the approximate molecular weight of β-arrestin. Since β-arrestin has been shown to be regulated by S-nitrosylation ^24,31^, we next performed a histological analysis of β-arrestin1/2 S-nitrosylation to test whether the glucocorticoid-induced reduction in total S-nitrosylation is due to changing the nitrosylation state of β-arrestin1/2. We combined the biotin-switch nitrosylation assay with a proximity ligation assay using a biotin antibody and a β-arrestin1/2 antibody to detect nitrosylated β-arrestin. Proximity ligation assay reveals protein-protein interactions with a 40-nm resolution ^32^. Cells were analyzed for the total number of biotin-β-arrestin interactions (i.e., nitrosylation association with β-arrestin) and for nuclear versus cytosolic localization of the interactions. Treatments with NE (1 µM), CORT (2 µM), and NE + CORT all failed to produce a statistically significant change in the PLA signal with respect to vehicle, suggesting they did not affect the nitrosylation of β-arrestin (Fig. 6F, G). The NE + CORT treatment caused a small decrease in the PLA signal, but this did not reach statistical significance (p = 0.1470, Dunnett’s multiple comparisons test). Similarly, the treatments failed to change the nuclear fraction of nitrosylated β-arrestin (Fig. 6H). These findings suggest that the nitrosylation dependence of the NE and Cort effects on ARa1b trafficking is not mediated by the regulation of β-arrestin1/2 S-nitrosylation and that nitrosylated β-arrestin1/2 is not trafficked to the nucleus in response to NE and/or Cort.

Ubiquitination is critical for endosomal processing and cellular trafficking. If glucocorticoid changes the internal trafficking of ARα1b, then it may do so by changing the ubiquitination state of β-arrestin ^33^. We tested for changes in the ubiquitination of β-arrestin in response to NE and CORT treatment (20 min). Both NE (1 µM) and CORT (2 µM) alone increased the ubiquitination of immunoprecipitated β-arrestin1/2 approximately two-fold (Supplemental Fig. 2). The increases in ubiquitinated β-arrestin1/2 caused by CORT and NE were additive when the cells were co-treated with the two drugs (Supplemental Fig. 2). Since the ubiquitination pattern of β-arrestin has been shown to affect trafficking of internalized cargo ^34^, these data suggest that NE and CORT may alter ARα1b receptor trafficking by increasing β-arrestin ubiquitination.

### Glucocorticoid receptor interaction with the **α**1 adrenoreceptor

To interrogate the mechanism by which NE and CORT exert cooperative effects on ARα1b trafficking and β-arrestin ubiquitination, we tested for a direct interaction between ARα1b and the glucocorticoid receptor (GR) with the proximity ligation assay. A human ARα1b construct with a myc-DDK tag (hARα1b-myc-DDK) or an ARα1b-eGFP construct was transiently transfected into N42 cells. Proximity ligation was achieved by using either a mouse anti-myc antibody or a mouse anti-GFP antibody to target ARα1b and a rabbit anti-glucocorticoid receptor antibody (M-20, Santa Cruz) to target GR. Individual cells were analyzed for the number of protein-protein interactions and for nuclear versus cytosolic localization of the interacting proteins. A basal level of interaction between ARα1b and GR was observed in vehicle-treated cells (Fig. 7A). Norepinephrine alone had no effect on ARα1b-GR interaction, whereas CORT treatment caused a small but significant decrease in total ARα1b-GR interaction compared to vehicle and NE treatment, which was unchanged by the combination of CORT + NE (Fig. 7A, B). We next analyzed the nuclear translocation of the ARα1b-GR complex after treatment with NE and/or CORT. Norepinephrine alone had no effect on the nuclear fraction of the ARα1b-GR complex, but CORT caused a significant increase in the nuclear ARα1b-GR PLA signal compared to vehicle and to NE. The increase in nuclear localization of the ARa1b-GR complex caused by CORT was reversed by co-application of NE (Fig. 7A, C). This suggested that NE activation of ARα1b prevented the CORT-induced nuclear translocation of the GR-ARα1b complex. Thus, ARα1b and GR can be found in a protein complex that is trafficked to the nucleus in response to CORT but not in the presence of NE, when the ARα1b is presumably mostly bound to its ligand.

**Figure 7.**
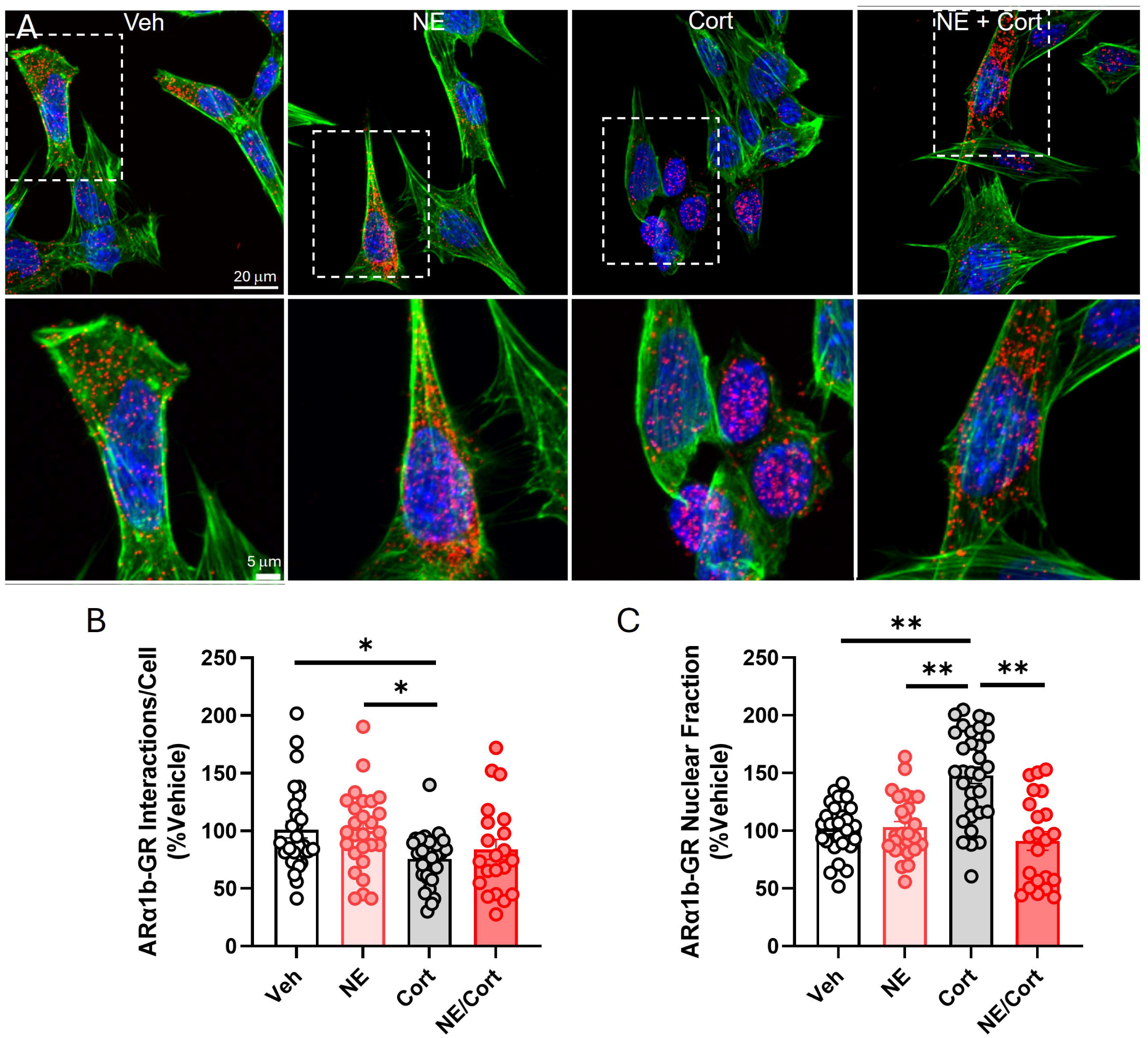
Interaction between ARα1b and GR. **A.** Representative images of ARα1b-myc-DDK-transfected N42 cells after the different treatments and following PLA. Lower images are high-magnification of boxes in upper images. Red = PLA signal, blue = DAPI staining of nuclei, green = phalloidin-FITC staining of F-actin **B.** Summary graph of total numbers of ARα1b-GR interactions under the different drug conditions. There was an overall decrease in interactions with Cort treatment [F(3, 102) = 3.831 P=0.012]. **C.** Summary graph of the nuclear fraction of ARα1b-GR profiles under the different treatments. Cort treatment significantly increased nuclear localization of the receptor complex relative to vehicle and the other treatment groups [F (3, 103) = 17.30 P<0.0001]. n = 21-31 cells from 3 separate assays. Data were analyzed by one-way ANOVA followed by Tukey’s multiple comparisons test. * p<0.05, ** p<0.0001.

### **β**-arrestin interactions with the **α**1 adrenoreceptor and glucocorticoid receptor

We next examined interactions of β-arrestin1/2 to determine whether the post-translational S-nitrosylation of β-arrestin is accompanied by changes in its interactions with ARα1b and GR. As expected, NE treatment (1 μM) of the N42 cells for 20 min caused an increase in ARα1b-β-arrestin interaction (Fig. 8A, B). Treatment with CORT alone (2 μM) had no effect on the number of ARα1b-β-arrestin PLA profiles, but it blocked the NE-induced increase in ARα1b-β-arrestin complex formation when co-applied with NE (Fig. 8A, B). CORT treatment alone also caused a significant increase in nuclear localization of the ARα1b/β-arrestin complex, which was blocked by NE when NE and CORT were co-applied (Fig. 8A, C).

**Figure 8.**
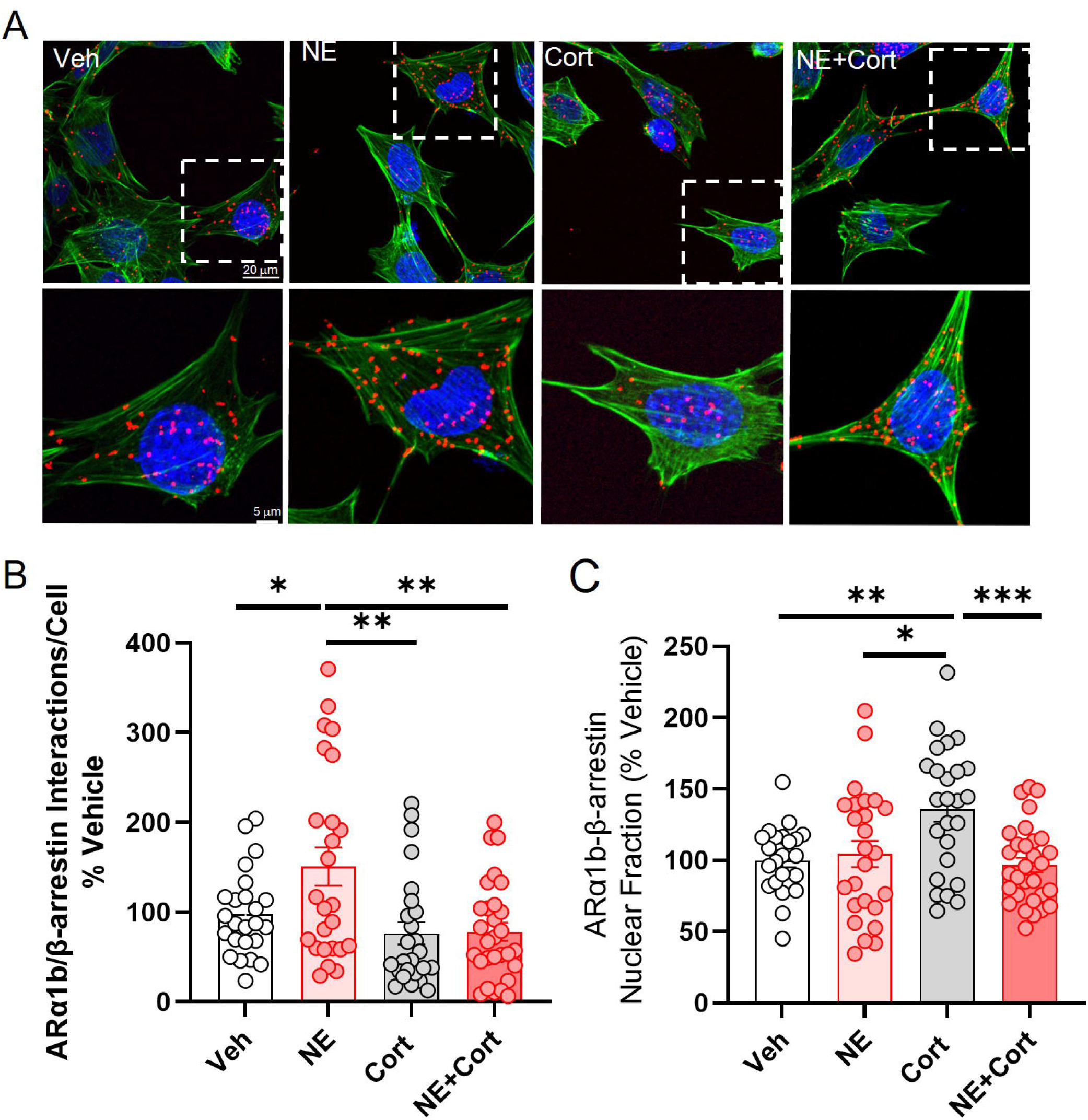
Interaction between ARα1b and β-arrestin. Proximity ligation assay with antibodies to tagged ARα1b and β-arrestin1/2. N42 cells were treated with drug for 20 min, followed by proximity ligation assay. **A.** Representative images. Blue = DAPI, red = PLA signal, green = phalloidin-FITC stain of F-actin. **B.** Treatments with NE, Cort, and NE+Cort. NE increased the number of ARa1b-b-arrestin interactions per cell; Cort alone had no effect, but blocked the NE-induced increase in ARα1b-β-arrestin interactions [F(3,99), p = 6.157, p = 0.0007], **C.** Cort treatment increased the nuclear translocation of the ARα1b-β-arrestin1/2 complex [F (3, 99) = 6.314, p = 0.0006], n = 23-30 cells from 3 separate assays. * p < 0.05. ** p<0.01, ***p<0.001. Data were analyzed by one-way ANOVA Followed by Tukey’s multiple comparisons test. Scale bar = 20 µm (main), 5 µm (inset).

Similar to what has been reported previously ^35^, we found a baseline interaction between GR and β-arrestin1/2. The drug treatments did not affect the total number of GR-β-arrestin1/2 interactions (Fig. 9A, B). CORT application caused the GR-β-arrestin1/2 complex to translocate to the nucleus, and co-application of NE with CORT did not prevent the CORT-induced nuclear translocation of the GR-β-arrestin1/2 complex (Fig. 9A, C).

**Figure 9.**
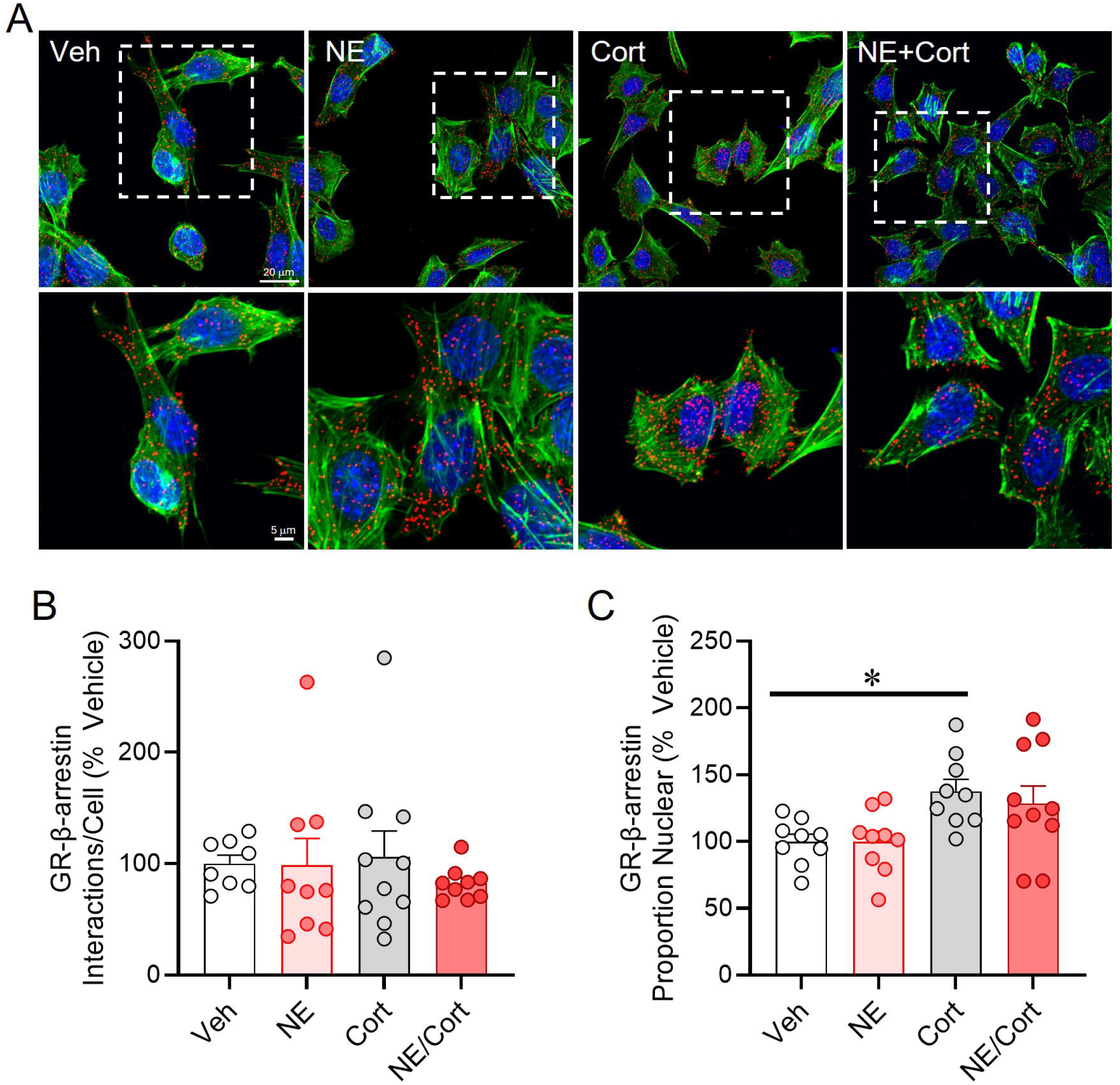
Interaction between GR and β-arrestin. Proximity ligation assay with antibodies to GR and β-arrestin1/2. **A.** Representative images of the PLA labeling with vehicle (Veh), NE, Cort, and NE + Cort. Lower images are high magnification of boxes in upper images. Blue = DAPI, red = PLA signal, green = phalloidin-FITC stain of F-actin. **B.** None of the treatments had an effect on the total number of GR-β-arrestin1/2 interactions per cell (one-way ANOVA, [F(3, 32) = 0.3233, p = 0.8085]. **C.** Cort treatment increased the nuclear fraction of the GR-β-arrestin1/2 complex (one-way ANOVA followed by Dunnett’s multiple comparisons test, [F(3, 33) = 4.058, p = 0.0147], n = 9-10 cells from 3 separate assays, * p < 0.05).

## Discussion

Acute stress desensitizes PVN CRH neurons in male mice to stimulation by noradrenergic afferents via a rapid glucocorticoid signaling mechanism. The glucocorticoid-induced NE desensitization suppresses the HPA response to somatic, but not psychological, stress and is dependent on the NE-mediated endocytosis of α1 adrenoreceptors ^21^. Here, we tested for the cellular mechanisms of the α1 adrenoreceptor desensitization in slices from male mice (consistent with our previous studies, Chen et al., 2019; Jiang et al., 2022), and in the N42 hypothalamic cell line, which expresses both CRH ^26^ and a putative membrane receptor that mediates rapid glucocorticoid signaling^21,36,37^.

Our findings reveal a rapid glucocorticoid regulation of the trafficking of α1 adrenoreceptors that diverts the receptors away from the rapid recycling endosomal pathway and into late endosomes. The glucocorticoid regulation of α1 receptor trafficking requires internalization of the α1 receptors by NE, is dependent on the overall nitrosylation state of the cells and involves ubiquitination of β-arrestin. The rapid glucocorticoid-induced re-routing of the α1 adrenoreceptors out of the rapid recycling endosomal pathway would be expected to deplete the cell of surface receptors and to cause the desensitization of CRH neurons to NE following acute stress exposure ^21^.

Glucocorticoid secretion after immune activation of the HPA axis is essential for downregulation of the immune response ^38^. Left unchecked, inflammation and hypotension associated with immune system activation can cause problems beyond the immune challenge itself. One example is toxic shock syndrome, which occurs when bacteria-produced toxins cause over-activation of cytokines and inflammatory cells resulting in fever, rash, hypotension, and, in extreme cases, organ failure and death^39^. Glucocorticoids constrain the immune response, and glucocorticoid therapy is often employed to inhibit the overactive immune system ^40^. For example, the cytokine storm occurring in some COVID-19 patients was treated with the synthetic glucocorticoid dexamethasone ^41,42^.

Previously we reported that both acute restraint stress and CORT pre-exposure induced a desensitization of CRH neurons to the excitatory effect of NE that lasted for the entire period of brain slice recording (2-6 h) ^21^. However, whether ex vivo brain slice preparation prevented the CRH neurons from recovering from their desensitization to NE by placing them in a suspended desensitized state was not known, nor was it clear how long the desensitization to NE lasts following stress. Here we sacrificed the mice at two time points following a 30-min restraint, at 4 h and 18 h. A 4-h interval falls within the normal recording period following slice preparation and, we reasoned, would determine whether the observed desensitization of the CRH neurons sustained *ex vivo* is an accurate reflection of what occurs *in vivo*. The 18-h interval fell on the day after the stress exposure and tested whether the *in vivo* duration of the NE desensitization was sustained overnight. We found that the 4-h post-stress recovery period was insufficient for the CRH neurons to recover from glucocorticoid-induced NE desensitization, and that the NE response was partially, though not entirely, restored at 18 h after the acute stress. Thus, the desensitization of CRH neurons to noradrenergic activation is long-lasting, persisting for over 18 h.

We showed previously that the CRH neuron response and HPA activation to a somatic stress, LPS exposure, but not to a psychological stress, predator odor, are mediated by NE afferents and susceptible to desensitization by prior stress exposure ^21^. The long-lasting desensitization to NE appears, therefore, to be a negative feedback filter of the HPA axis that passes acute psychological stress activation while filtering out more long-lasting somatic stress activation (e.g., by immune challenge). This filtering of stress modality-specific stress responsiveness is adaptive in that it reduces the duration of stress exposure, since somatic stressors tend to be prolonged in nature, which would otherwise subject the organism to the damaging effects of sustained glucocorticoid exposure. However, this feature could be maladaptive in modern day, since deprioritizing somatic stressors in favor of psychological stressors may be beneficial in life-or-death situations, such as a looming predator, but those situations are less likely to occur in modern human life. Additionally, many modern stressors could be perceived as severe even though they may not be life threatening. Thus, acute stress-induced desensitization could be a maladaptive mechanism that needlessly deprioritizes somatic stressors and leaves the body susceptible to physiological distress. Further studies are needed to determine whether desensitization of the HPA axis generalizes to other somatic stressors, and to ascertain the role of noradrenergic desensitization of the HPA axis in diseases known to be caused by HPA dysregulation.

### Glucocorticoid altered AR**α**1b trafficking

Live-cell imaging of NE and CORT effects on trafficking of ARα1b revealed that much of the trafficked receptor was directed to a single area of each cell, or a “hot spot”. This receptor-concentrated “hot-spot” co-localized well with the Rab7 marker of late endosomes. FRET analysis confirmed that Rab7 and ARα1b are physically associated within 10 nm of each other, indicating that ARα1b is likely located in the late endosome. The lack of increase in early endosomal ARα1b in the presence of CORT indicates that the desensitization to NE by CORT is not caused by an increase in ARα1b internalization, as we originally hypothesized based on its prevention by blocking dynamin-dependent endocytosis ^21^. The decrease in ARα1b in rapidly recycling endosomes suggests that the increase in cytosolic receptor is due to a decrease in receptor trafficking back to the membrane. This rapid steroid-driven change in GPCR trafficking is not only novel *per se*, but also reveals a previously unobserved switch in the speed of receptor trafficking. Receptors of the Class A GPCRs, including the adrenoreceptors, are rapidly recycled back to the plasma membrane in minutes, whereas Class B receptors, which bind more stably to β-arrestin, take several hours ^43^. The ubiquitination patterns of the β-arrestin that associates with these receptors during endocytosis appear to be specific for the receptor family type and ultimately control the speed of trafficking ^34^. Our findings reveal a previously unobserved switch in GPCR trafficking kinetics and suggest that CORT could be altering the ubiquitination patterns of β-arrestin to switch GPCRs from rapidly trafficking Class A receptors to slowly trafficking Class B-like receptors. The overall increase in β-arrestin ubiquitination in the presence of CORT suggests that other rapidly trafficking GPCRs may be similarly affected. Thus, this may represent a more universal mechanism of CORT desensitization of multiple neurotransmitters systems, a prospect that needs to be further explored.

While it remains unclear how glucocorticoids cause an increase in β-arrestin ubiquitination, changes in trafficking may also be achieved by S-nitrosylation of trafficking proteins. We found no change in the nitrosylation of β-arrestin 1/2 with NE or CORT treatment, indicating that the convergent regulation of ARα1 by nitrosylation and glucocorticoid is not mediated by a change in the nitrosylation state of β-arrestin. S-nitrosylation regulates multiple other proteins involved in GPCR trafficking in addition to β-arrestin, including G protein-coupled receptor kinase 2, endophilin, and clathrin (Hayashi et al., 2018; Ozawa et al., 2008; Whalen et al., 2007). It is possible, therefore, that rapid CORT regulation of nitrosylation of these proteins may mediate the glucocorticoid modulation of ARα1b intracellular trafficking, a possibility that remains to be determined.

The nuclear glucocorticoid receptor is one possible receptor candidate for the rapid CORT actions in this pathway. However, one or more other membrane receptors may also be candidates for the rapid glucocorticoid effects. Many studies have provided evidence for the existence of a glucocorticoid receptor on the membrane that signals via G protein or tyrosine kinase activity (Di et al., 2003; Rhen and Cidlowski, 2005), and a recent study shows glucocorticoid binding to a membrane adhesion Gαi-coupled orphan receptor ^45^. Our PLA analysis indicated a close physical association (<40 nm) of GR with ARα1b, which suggests a possible direct interaction of the two receptors. We have found the M-20 GR antibody we used here to recognize proteins in the membrane, cytosolic, and nuclear fractions in Western blot analyses (Figure 7). The GR/ARα1b complex detected by PLA responded to CORT by translocating to the nucleus, an effect that was blocked by concomitant activation of ARα1b by NE. Thus, the receptor complex is differentially regulated depending on its component receptor activation. The interaction between the two receptors suggests this as a possible rapid mechanism whereby glucocorticoids could influence ARα1b intracellular trafficking.

Based on the well-established role of β-arrestin in receptor trafficking, our finding of an additive effect of Cort and NE on β-arrestin ubiquitination suggests that β-arrestin is a locus of the Cort actions that regulate ARα1b trafficking. The proximity ligation assay showed that both GR and ARα1b can individually interact with β-arrestin, and further showed that GR and ARα1b can interact with each other. However, the PLA did not allow for determination of whether GR, ARα1b, and β-arrestin act together in a single complex. The ability of CORT to cause nuclear translocation of the ARα1b-β-arrestin complex, the ARa1b-GR complex, and the GR-β-arrestin complex suggests a trimeric complex that includes the GR, but we have not demonstrated the multimeric association of the three proteins by molecular assays. While adrenoreceptor localization in the nucleus has been previously reported in cardiomyocytes ^46^ and astrocytes ^47^, this phenomenon is still relatively novel. ARα1b signaling could be further biased upon entering the nucleus, giving rise to more potential targets for synergistic NE-CORT signaling. Regardless, co-activation of ARα1b and GR results in fundamental changes in the state of β-arrestin, increasing its ubiquitination, which are associated with profound alterations in the trafficking of the ARα1b. The CORT regulation of adrenoreceptor trafficking is likely to be responsible for the stress desensitization of the CRH neurons to the noradrenergic excitatory synaptic input activated by somatic stress.

Finally, the concept of CORT-induced changes in endosomal trafficking is novel. Changes in receptor trafficking could have a broader impact on neuronal function including plasticity via regulation of ionotropic receptors ^48,49^. Glucocorticoids have been shown to facilitate Rab4 cycling, in turn increasing AMPAR recycling ^50^. These studies are crucial for understanding the full extent of rapid stress-induced glucocorticoid regulation of synaptic signaling beyond the classically studied transcriptional regulation.

## Supporting information

Table 1. Antibodies used

## Acknowledgments

We would like to thank Dr. Chris Hague of the University of Washington for his donation of the ARa1b-eGFP plasmid. This work was supported by NIH grant MH119283 and a Carol Lavin Bernick Faculty Grant from Tulane University.

## Author contributions

Conceptualization: G.L.W, L.M.H., and J.G.T.; methodology: G.L.W., L.M.H, Z.J., and J.G.T.; investigation: G.L.W., L.M.H, Z.J., A.M.N., M.S.F., S.N., P.S.T.; writing – original draft: G.L.W. and L.M.H.; writing –review & editing: J.G.T., G.L.W., L.M.H., Z.J., and G.L.W.; visualization: G.L.W., L.M.H., Z.J., and J.G.T.; supervision, J.G.T.; project administration, J.G.T.; funding acquisition, J.G.T.

## Supplemental information

Document S1. Figures S1-S3

Video S1

## Methods

### Mice

CRH-eGFP transgenic mice were raised in-house from breeders provided by the Mutant Mouse Resource and Research Center at the University of California, Davis (MMRRC, stock: Tg(Crh-eGFP)HS57Gsat/Mm, RRID:MMRRC_017058-UCD). The mice were genotyped at 2-3 weeks using the primer set: CRH-F1 (CTG TCT TGT CGT GGG TGT CCG AT); GFP R2 (TAG CGG CTG AAG CAC TGC A). All animals and procedures were approved by the Tulane Institutional Animal Care and Use Committee (IACUC) and followed National Institutes of Health guidelines. Mice were housed in an AAALAC-accredited animal facility on a 12:12 light/dark cycle (lights on at 7:00 AM) under controlled temperature (20°C) and received food and water *ad libitum*.

### Cell culture

We used an immortalized hypothalamic cell line, mHYPOE-N42 (N42) cells (Cellutions Biosystems), that expresses CRH ^26^ as well as both the nuclear glucocorticoid receptor and a membrane-associated glucocorticoid receptor ^36,37^. Cells were plated at a density of 4.5 x 10^4^ cells/well on 12 mm glass coverslips (size 0, Carolina Biological, Burlington, NC) in 24-well plates (Corning, Corning, NY) in 1X Dulbecco’s Modified Eagle’s Medium (DMEM, Millipore Sigma, Burlington, MA) supplemented with 10% fetal bovine serum (FBS, Atlas Biologicals), 1% penicillin-streptomycin solution, and 0.2% Plasmocin Prophylactic® (Invivogen, San Diego, CA). Plates were incubated at 37°C with 5% CO_2_ until cells were approximately 80% confluent. To remove endogenous hormones, the culture medium was then removed, cells were washed once with 1X Dulbecco’s-phosphate buffered saline (PBS), and fresh DMEM without phenol red supplemented with 5% charcoal-stripped (CS) FBS was added for 16 h at 37°C with 5% CO_2_. For proximity ligation assays, N42 cells were seeded at a density of 12,000/well in 16-well chamber slides and cultured as described above.

### Alpha-1 adrenoreceptor overexpression

For live imaging assays, N42 cells were transiently transfected with a pEGFP-ARα1b construct generously donated by Dr. Chris Hague of the University of Washington. Transfection was achieved by electroporation with two 10 ms, 1400 mV pulses using the NEON® Transfection System (Life Technologies, Carlsbad, CA). Transfected cells were plated onto No. 1 coverslip bottom plates for live-cell imaging with a Nikon A1 confocal microscope with incubator stage (Nikon USA, Melville, NY). For proximity ligation assays, cells were transfected with either pEGFP-ARα1b or Myc-DDK-tagged human ARα1b (Origene, Rockville, MD) using Lipofectamine 3000.

### Cell imaging

Intracellular ARα1b trafficking was tracked in live N42 cells using cytosolic intensity analysis, Förster resonance energy transfer (FRET), or pixel-based colocalization. For cytosolic intensity analysis, cells were transiently transfected as described above with pEGFP-ARα1b. Images were captured every 15 seconds and intensity was measured within a manually defined cytosolic region of interest. For FRET, cells were transiently transfected with a combination of pEGFP-ARα1b (the FRET donor) and Rab4a-dsRed, Rab5-dsRed, Rab7-dsRed, or Rab11-dsRed (FRET acceptors) and imaged repeatedly over time. Three images were taken at each time point: 1) donor excitation, donor emission, 2) acceptor excitation, acceptor emission, and 3) donor excitation, acceptor emission. The same set of images was also taken in cells only expressing either donor or acceptor fluorophores and used as bleed-through controls. Images were analyzed using the FRET analysis tool in Image J ^51^, which yielded images with a FRET intensity for each pixel. A region of interest around each cell expressing both constructs was defined, and the average FRET signal was calculated in the region to quantify the colocalization of ARα1b with each Rab protein. For pixel-based colocalization, cells were transfected with pEGFP-ARα1b and RFP-LAMP1 followed by colocalization analysis using Cell Profiler ^52^ to calculate each cell’s mean Pearson’s coefficient.

### Ubiquitination assay

N42 cells were treated with drugs for 20 min at 37°C before lysis with NP40 lysis buffer containing 50 mM Tris-HCl (pH 7.4), 150 mM NaCl, 1% NP-40 and 5 mM EDTA for 1 h at 4°C. Immunoprecipitation (IP) of the protein of interest was performed using the Catch-and-Release kit (Millipore Sigma, Burlington, MA), eluting with the non-denaturing buffer. Eluates were run on a gradient SDS-PAGE gel (Bio-Rad, Hercules, CA) and transferred to PVDF membrane. Blots were blocked for 1 h in 5% dry milk in Tris-buffered saline with 0.05% tween (TBST). Primary antibodies were applied in 5% BSA in TBST overnight, followed by 4 washes in TSBT. HRP-conjugated secondary antibody (Clean-Blot IP Detection Reagent, Thermo-Fisher, Waltham MA) was applied in 5% dry milk in TBST for 1 h at room temperature followed by 4 washes in TBST. Chemiluminescent substrate (Super Signal West Pico, Bio-Rad, Hercules, CA) was applied for 5 min followed by detection on a ChemiDoc densitometer (Bio-Rad). Densitometry scores for ubiquitin were quantified and normalized against the protein purified by immunoprecipitation. The antibodies used to target GFP, β-arrestin, and ubiquitin are described in Table 1.

### Proximity ligation Assay

Proximity ligation assay (PLA) was carried out with the Duolink system (Millipore Sigma). N42 cells were seeded at a density of 12,000 cells/well on chamber slides (Thermo Scientific) and cultured as described above. For some assays, cells were transfected with human ARα1b-myc-DDK (Origene) or pEGFP-ARα1b using Lipofectamine 3000 (Thermo Scientific). Before assay, cells were cultured for 16h in phenol red-free DMEM supplemented with 10% CS FBS, pen-strep, and Plasmocin. Cells were treated for 20 min with the indicated drugs at 37°C, followed by three washes with PBS, fixation in 4% paraformaldehyde, and blocking with Duolink blocking solution. Incubation in primary antibody was carried out overnight at 4°C. For each assay, two primary antibodies were used, one for each protein or post-translational modification. One antibody of each set was prepared in rabbit and the other in mouse in order to use the complementary secondary antibodies of the Duolink system. After washing with PBS, PLA was performed with the Duolink In Situ system according to the manufacturer’s instructions (Millipore Sigma, Burlington, MA). Negative control wells were incubated with only one of the two primary antibodies and generally yielded less than 5 signals per cell (Supplemental Figure 3). For some assays, PLA was followed by incubation in phalloidin-FITC (1:750, Abcam, Boston, MA) to demarcate cells. Cells were imaged at 40x or 60x with a Nikon Eclipse Ti confocal microscope. For each treatment group, Z-stacks were taken from 2-4 fields. Image quantification was performed with NIH ImageJ/FIJI software. Protein-protein interaction signal was quantified as the number of “dots”. For cells expressing the ARα1b-myc-DDK transgene, a threshold of 15 fluorescent dots per cell was used to distinguish the stain from background, and the phalloidin-FITC stain was used to demarcate the cell boundary. For assays employing pEGFP-ARα1b, transfected cells were determined by GFP signal, and these cells were individually analyzed for total number and nuclear fraction of interactions. See Table 1 for antibodies and dilutions.

### Nitrosylation assay

For histological nitrosylation assays, cells were cultured on chamber slides as described above. A biotin switch assay was performed according to the manufacturer’s instructions (Cayman Chemical Company, Ann Arbor, MI) to determine nitrosylation of endogenous β-arrestin1/2. After substitution of biotin for the NO group, PLA was performed with mouse anti-biotin and rabbit anti-β-arrestin1/2 antibodies (see Table 1 for antibody information). Phalloidin-FITC staining was not compatible with the biotin switch assay, so entire fields, rather than individual cells, were quantified for the post-translational modification signal. For each field, the total number of nuclei and the total number of PLA signals were determined in their individual channels. An average number of PLA (nitrosylation) signals per cell was determined by dividing the total number of PLA signals by the total number of nuclei. The nuclear outlines were superimposed onto the image of PLA signal, the PLA signal outside the nuclei was cleared, and the remaining PLA signal was determined to calculate the average nuclear fraction for each field.

For assays of nitrosylation in cell lysates, N42 cells were seeded at 0.5 x 10^6^ in 6-well culture plates and grown to confluence. After treatment for 20 min with the indicated drugs at 37°C, cells were lysed, and NO post-translational modifications were substituted with a tandem mass tag (TMT) according to the manufacturer’s instructions (Thermo Pierce). Lysates were subjected to Western blotting as described above and probed with an anti-TMT antibody. Densitometry was performed with Image Lab software (Bio-Rad), and results are presented as the sum density of all bands in each lane.

### Electrophysiology

Ex vivo whole-cell patch clamp electrophysiological recordings were conducted in acutely prepared hypothalamic slices from 6-9 week-old male CRH-eGFP mice. On the mornings of experiments, a mouse was gently removed from its home cage to a transfer cage and transported to an adjacent room in the vivarium, where it was immobilized in a flexible plastic decapitation cone (DecapiCone, Braintree Scientific) and, within less than 2 min of removal from its home cage, decapitated without anesthesia using a rodent guillotine. Anesthetics activate the HPA axis and increase circulating levels of ACTH and CORT ^53^, which otherwise remain low in our hands for ∼3 min from the start of handling. Following decapitation, the brain was quickly removed and cooled in oxygenated, ice-cold artificial cerebrospinal fluid (aCSF) containing (in mM): 140 NaCl, 3 KCl, 1.3 MgSO_4_, 11 Glucose, 5 HEPES, 1.4 NaH2PO_4_, 3.25 NaOH, 2.4 CaCl_2_, pH 7.2-7.4, with an osmolarity of 290-300 mOsm. The forebrain was isolated, the ventral half was blocked, and the caudal surface of the block was glued to a vibratome chuck. Two or three 300 μm-thick coronal slices containing the hypothalamic PVN were then sectioned on a vibrating slicer (Leica) in cooled aCSF, bisected down the midline, and the hemi-slices were transferred to a holding chamber, where they were maintained in oxygenated aCSF at 20°C for at least 1 h to allow for recovery prior to electrophysiological recordings. Single slices were then transferred and weighted with a silver wire to the bottom of a submerged recording chamber on a fixed-stage, upright microscope (Olympus BXW51) equipped with a long working distance, water immersion 40x objective, where they were perfused with fresh aCSF at a rate of ∼2 ml/min at 20°C. eGFP-expressing CRH neurons in the PVN were first identified under epifluorescence illumination, which was then switched to infrared-differential interference contrast optics and the cells were visualized on a monitor using a near-infrared camera to target them for whole-cell patch clamp recordings. Patch pipettes were pulled from borosilicate glass (ID 1.2 mm, OD 1.65 mm) to a resistance of 3-6 MΩ on a horizontal puller (P-97, Sutter Instr.) and filled with an internal patch solution containing (in mM): 120 potassium gluconate, 10 KCl, 1 NaCl, 1 MgCl_2_, 0.1 CaCl_2_, 5.5 EGTA, 10 HEPES, 2 Mg-ATP, and 0.3 Na-GTP, with a pH adjusted to 7.3 with KOH and osmolarity adjusted to 300 mOsm with D-sorbitol. The GABA_A_ receptor antagonist picrotoxin (PTX, 50 µM) was applied via the bath perfusion to isolate excitatory postsynaptic currents (EPSCs). While multiple cells from the same mouse were sometimes recorded, only one cell was recorded per hemi-slice. The numbers of recorded CRH neurons are designated as the ‘n’ and numbers of mice used for each experiment are designated as the ‘N’.

### Statistical analyses

Data are presented as the mean ± standard error of the mean. Postsynaptic currents were selected and analyzed for changes in frequency, amplitude, and decay time with Minianalysis 6.0 (Synaptosoft Inc.). Baseline values were calculated from 3 min of recording just prior to drug application and compared with the 3 min of recording at the peak of drug responses using a within-cell analysis. For live cell imaging experiments, mean fluorescence values were calculated from the final 3 min of the baseline and of the drug application and were compared using a within-cell analysis. FRET values were taken from single time points at the end of both the baseline and drug applications. Statistical significance was determined with the two-tailed, paired Student’s *t*-test for within-cell drug effects, or ANOVA with post-hoc Dunnett’s or Tukey’s analysis where indicated (GraphPad Prism 7-9 and SigmaPlot 11.0, Systat Software, Inc.).

## References

1. Kuo, T., McQueen, A., Chen, T.-C., and Wang, J.-C. (2015). Regulation of Glucose Homeostasis by Glucocorticoids. In Advances in experimental medicine and biology, pp. 99–126. 10.1007/978-1-4939-2895-8_5.

2. Oppong, E., and Cato, A.C.B. (2015). Effects of Glucocorticoids in the Immune System. In Advances in experimental medicine and biology, pp. 217–233. 10.1007/978-1-4939-2895-8_9.

3. Whitnall, M.H. (1993). Regulation of the hypothalamic corticotropin-releasing hormone neurosecretory system. Prog Neurobiol 40, 573–629. 10.1016/0301-0082(93)90035-Q.

4. Pratt, W.B., Morishima, Y., Murphy, M., and Harrell, M. (2006). Chaperoning of glucocorticoid receptors. Handb Exp Pharmacol, 111–138.

5. Lightman, S.L., and Young, W.S. (1989). Influence of steroids on the hypothalamic corticotropin-releasing factor and preproenkephalin mRNA responses to stress. Proc Natl Acad Sci U S A 86, 4306– 4310.

6. Di, S., Malcher-Lopes, R., Halmos, K.C., and Tasker, J.G. (2003). Nongenomic glucocorticoid inhibition via endocannabinoid release in the hypothalamus: a fast feedback mechanism. J Neurosci 23, 4850–4857.

7. Tasker, J.G., Di, S., and Malcher-Lopes, R. (2006). Minireview: rapid glucocorticoid signaling via membrane-associated receptors. Endocrinology 147, 5549–5556. 10.1210/en.2006-0981.

8. Evanson, N.K., Tasker, J.G., Hill, M.N., Hillard, C.J., and Herman, J.P. (2010). Fast Feedback Inhibition of the HPA Axis by Glucocorticoids Is Mediated by Endocannabinoid Signaling. Endocrinology 151, 4811–4819. 10.1210/en.2010-0285.

9. Gjerstad, J.K., Lightman, S.L., and Spiga, F. (2018). Role of glucocorticoid negative feedback in the regulation of HPA axis pulsatility. Stress 21, 403–416. 10.1080/10253890.2018.1470238.

10. Evanson, N.K., Tasker, J.G., Hill, M.N., Hillard, C.J., and Herman, J.P. (2010). Fast feedback inhibition of the HPA axis by glucocorticoids is mediated by endocannabinoid signaling. Endocrinology 151, 4811–4819. 10.1210/en.2010-0285.

11. Cole, R.L., and Sawchenko, P.E. (2002). Neurotransmitter regulation of cellular activation and neuropeptide gene expression in the paraventricular nucleus of the hypothalamus. J Neurosci 22, 959–969.

12. Domyancic, A. V, and Morilak, D.A. (1997). Distribution of 1A Adrenergic Receptor mRNA in the Rat Brain Visualized by In Situ Hybridization. J. Comp. Neurol 386, 358–378. 10.1002/(SICI)1096-9861(19970929)386:3.

13. Williams, A.M., and Morilak, D.A. (1996). α(1B) adrenoceptors in rat paraventricular nucleus overlap with, but do not mediate, the induction of c-Fos expression by osmotic or restraint stress. Neuroscience 76, 901–913. 10.1016/S0306-4522(96)00351-X.

14. Sands, S.A., and Morilak, D.A. (1999). Expression of alpha1D adrenergic receptor messenger RNA in oxytocin- and corticotropin-releasing hormone-synthesizing neurons in the rat paraventricular nucleus. Neuroscience 91, 639–649. 10.1016/S0306-4522(98)00616-2.

15. Chen, C., Jiang, Z.Y., Fu, X., Yu, D., Huang, H., and Tasker, J.G. (2019). Astrocytes Amplify Neuronal Dendritic Volume Transmission Stimulated by Norepinephrine. Cell Rep 29, 4349. 10.1016/J.CELREP.2019.11.092.

16. Jiang, Z., Chen, C., Weiss, G.L., Fu, X., Stelly, C.E., Sweeten, B.L.W., Tirrell, P.S., Pursell, I., Stevens, C.R., Fisher, M.O., et al. (2022). Stress-induced glucocorticoid desensitizes adrenoreceptors to gate the neuroendocrine response to somatic stress in male mice. Cell Rep 41, 111509. 10.1016/J.CELREP.2022.111509.

17. Ericsson, a, Kovács, K.J., and Sawchenko, P.E. (1994). A functional anatomical analysis of central pathways subserving the effects of interleukin-1 on stress-related neuroendocrine neurons. J Neurosci 14, 897–913.

18. Herman, J.P., Figueiredo, H., Mueller, N.K., Ulrich-Lai, Y., Ostrander, M.M., Choi, D.C., and Cullinan, W.E. (2003). Central mechanisms of stress integration: hierarchical circuitry controlling hypothalamo-pituitary-adrenocortical responsiveness. Front Neuroendocrinol 24, 151–180.

19. Takemura, T., Makino, S., Takao, T., Asaba, K., Suemaru, S., and Hashimoto, K. (1997). Hypothalamic-pituitary-adrenocortical responses to single vs. repeated endotoxin lipopolysaccharide administration in the rat. Brain Res 767, 181–191. 10.1016/S0006-8993(97)00460-5.

20. Bienkowski, M.S., and Rinaman, L. (2008). Noradrenergic inputs to the paraventricular hypothalamus contribute to HPA axis and central Fos activation in rats after acute systemic endotoxin exposure. Neuroscience 156, 1093. 10.1016/J.NEUROSCIENCE.2008.08.011.

21. Jiang, Z., Chen, C., Weiss, G.L., Fu, X., Stelly, C.E., Sweeten, B.L.W., Tirrell, P.S., Pursell, I., Stevens, C.R., Fisher, M.O., et al. (2022). Stress-induced glucocorticoid desensitizes adrenoreceptors to gate the neuroendocrine response to somatic stress in male mice. Cell Rep 41. 10.1016/J.CELREP.2022.111509.

22. Claing, A., Laporte, S.A., Caron, M.G., and Lefkowitz, R.J. (2002). Endocytosis of G protein-coupled receptors: roles of G protein-coupled receptor kinases and ß-arrestin proteins. Prog Neurobiol 66, 61–79. 10.1016/S0301-0082(01)00023-5.

23. Shenoy, S.K., Modi, A.S., Shukla, A.K., Xiao, K., Berthouze, M., Ahn, S., Wilkinson, K.D., Miller, W.E., and Lefkowitz, R.J. (2009). -Arrestin-dependent signaling and trafficking of 7-transmembrane receptors is reciprocally regulated by the deubiquitinase USP33 and the E3 ligase Mdm2. Proceedings of the National Academy of Sciences 106, 6650–6655. 10.1073/pnas.0901083106.

24. Ozawa, K., Whalen, E.J., Nelson, C.D., Mu, Y., Hess, D.T., Lefkowitz, R.J., and Stamler, J.S. (2008). S-nitrosylation of beta-arrestin regulates beta-adrenergic receptor trafficking. Mol Cell 31, 395–405. 10.1016/j.molcel.2008.05.024.

25. Day, H.E.W., Campeau, S., Watson, S.J., and Akil, H. (1999). Expression of α1b Adrenoceptor mRNA in Corticotropin-Releasing Hormone-Containing Cells of the Rat Hypothalamus and Its Regulation by Corticosterone. The Journal of Neuroscience 19, 10098. 10.1523/JNEUROSCI.19-22-10098.1999.

26. Dalvi, P.S., Nazarians-Armavil, A., Tung, S., and Belsham, D.D. (2011). Immortalized neurons for the study of hypothalamic function. Am J Physiol Regul Integr Comp Physiol 300, 1030–1052. 10.1152/AJPREGU.00649.2010/ASSET/IMAGES/LARGE/ZH60041175260006.JPEG.

27. Stenmark, H. (2009). Rab GTPases as coordinators of vesicle traffic. Nat Rev Mol Cell Biol 10, 513–525. 10.1038/nrm2728.

28. Li, H., Li, H.-F., Felder, R.A., Periasamy, A., and Jose, P.A. (2008). Rab4 and Rab11 coordinately regulate the recycling of angiotensin II type I receptor as demonstrated by fluorescence resonance energy transfer microscopy. J Biomed Opt 13, 031206. 10.1117/1.2943286.

29. Whalen, E.J., Foster, M.W., Matsumoto, A., Ozawa, K., Violin, J.D., Que, L.G., Nelson, C.D., Benhar, M., Keys, J.R., Rockman, H.A., et al. (2007). Regulation of beta-adrenergic receptor signaling by S-nitrosylation of G-protein-coupled receptor kinase 2. Cell 129, 511–522. 10.1016/J.CELL.2007.02.046.

30. Jaffrey, S.R., Erdjument-Bromage, H., Ferris, C.D., Tempst, P., and Snyder, S.H. (2001). Protein S-nitrosylation: a physiological signal for neuronal nitric oxide. Nat Cell Biol 3, 193–197. 10.1038/35055104.

31. Hayashi, H., Hess, D.T., Zhang, R., Sugi, K., Gao, H., Tan, B.L., Bowles, D.E., Milano, C.A., Jain, M.K., Koch, W.J., et al. (2018). S-Nitrosylation of β-Arrestins Biases Receptor Signaling and Confers Ligand Independence. Mol Cell 70, 473–487.e6. 10.1016/J.MOLCEL.2018.03.034.

32. Alam, M.S. (2018). Proximity Ligation Assay (PLA). Curr Protoc Immunol 123, e58. 10.1002/CPIM.58.

33. Patwardhan, A., Cheng, N., and Trejo, J. (2021). Post-Translational Modifications of G Protein– Coupled Receptors Control Cellular Signaling Dynamics in Space and Time. Pharmacol Rev 73, 120. 10.1124/PHARMREV.120.000082.

34. Shenoy, S.K., Barak, L.S., Xiao, K., Ahn, S., Berthouze, M., Shukla, A.K., Luttrell, L.M., and Lefkowitz, R.J. (2007). Ubiquitination of beta-Arrestin Links Seven-transmembrane Receptor Endocytosis and ERK Activation. Journal of Biological Chemistry 282, 29549–29562. 10.1074/jbc.M700852200.

35. Petrillo, M.G., Oakley, R.H., and Cidlowski, J.A. (2019). β-Arrestin-1 inhibits glucocorticoid receptor turnover and alters glucocorticoid signaling. J Biol Chem 294, 11225. 10.1074/JBC.RA118.007150.

36. Weiss, G.L., Rainville, J.R., Zhao, Q., and Tasker, J.G. (2019). Purity and Stability of the Membrane-Limited Glucocorticoid Receptor Agonist Dexamethasone-BSA. Steroids 142, 2. 10.1016/J.STEROIDS.2017.09.004.

37. Rainville, J.R., Weiss, G.L., Evanson, N., Herman, J.P., Vasudevan, N., and Tasker, J.G. (2019). Membrane-initiated nuclear trafficking of the glucocorticoid receptor in hypothalamic neurons. Steroids 142, 55–64. 10.1016/J.STEROIDS.2017.12.005.

38. Pace, T.W.W., and Miller, A.H. (2009). Cytokines and glucocorticoid receptor signaling. Relevance to major depression. Ann N Y Acad Sci 1179, 86–105. 10.1111/j.1749-6632.2009.04984.x.

39. Davis, J.P., Chesney, P.J., Wand, P.J., and LaVenture, M. (1980). Toxic-shock syndrome: epidemiologic features, recurrence, risk factors, and prevention. N Engl J Med 303, 1429–1435. 10.1056/NEJM198012183032501.

40. Rhen, T., and Cidlowski, J.A. (2005). Antiinflammatory Action of Glucocorticoids — New Mechanisms for Old Drugs. New England Journal of Medicine 353, 1711–1723. 10.1056/NEJMra050541.

41. Wagner, C., Griesel, M., Mikolajewska, A., Mueller, A., Nothacker, M., Kley, K., Metzendorf, M.I., Fischer, A.L., Kopp, M., Stegemann, M., et al. (2021). Systemic corticosteroids for the treatment of COVID-19. Cochrane Database Syst Rev 2021, 14963. 10.1002/14651858.CD014963.

42. Ahmed-Hassan, H., Sisson, B., Shukla, R.K., Wijewantha, Y., Funderburg, N.T., Li, Z., Hayes, D., Demberg, T., and Liyanage, N.P.M. (2020). Innate Immune Responses to Highly Pathogenic Coronaviruses and Other Significant Respiratory Viral Infections. Front Immunol 11. 10.3389/FIMMU.2020.01979.

43. Conway, B.R., Minor, L.K., Xu, J.Z., D’Andrea, M.R., Ghosh, R.N., and Demarest, K.T. (2001). Quantitative analysis of agonist-dependent parathyroid hormone receptor trafficking in whole cells using a functional green fluorescent protein conjugate. J Cell Physiol 189, 341–355. 10.1002/jcp.10028.

44. Di, S., Malcher-Lopes, R., Halmos, K.C., and Tasker, J.G. (2003). Nongenomic glucocorticoid inhibition via endocannabinoid release in the hypothalamus: a fast feedback mechanism. J Neurosci 23, 4850–4857.

45. Ping, Y.Q., Mao, C., Xiao, P., Zhao, R.J., Jiang, Y., Yang, Z., An, W.T., Shen, D.D., Yang, F., Zhang, H., et al. (2021). Structures of the glucocorticoid-bound adhesion receptor GPR97–Go complex. Nature 2020 589:7843 589, 620–626. 10.1038/s41586-020-03083-w.

46. Wu, S.C., and Oconnell, T.D. (2015). Nuclear Compartmentalization of α1-Adrenergic Receptor Signaling in Adult Cardiac Myocytes. J Cardiovasc Pharmacol 65, 91. 10.1097/FJC.0000000000000165.

47. Benton, K.C., Wheeler, D.S., Kurtoglu, B., Ansari, M.B.Z., Cibich, D.P., Gonzalez, D.A., Herbst, M.R., Khursheed, S., Knorr, R.C., Lobner, D., et al. (2022). Norepinephrine activates β1-adrenergic receptors at the inner nuclear membrane in astrocytes. Glia 70, 1777–1794. 10.1002/GLIA.24219.

48. Groc, L., Choquet, D., and Chaouloff, F. (2008). The stress hormone corticosterone conditions AMPAR surface trafficking and synaptic potentiation. Nat Neurosci 11, 868–870. 10.1038/NN.2150.

49. Krugers, H.J., Hoogenraad, C.C., and Groc, L. (2010). Stress hormones and AMPA receptor trafficking in synaptic plasticity and memory. Nat Rev Neurosci 11, 675–681. 10.1038/NRN2913.

50. Liu, W., Yuen, E.Y., and Yan, Z. (2010). The Stress Hormone Corticosterone Increases Synaptic α-Amino-3-hydroxy-5-methyl-4-isoxazolepropionic Acid (AMPA) Receptors via Serum- and Glucocorticoid-inducible Kinase (SGK) Regulation of the GDI-Rab4 Complex. Journal of Biological Chemistry 285, 6101–6108. 10.1074/JBC.M109.050229.

51. Marchal, O., and Converset, N. (2006). FRET and Colocalization Analyzer. https://imagej.nih.gov/ij/plugins/fret-analyzer/fret-analyzer.htm.

52. Carpenter, A.E., Jones, T.R., Lamprecht, M.R., Clarke, C., Kang, I.H., Friman, O., Guertin, D. a, Chang, J.H., Lindquist, R. a, Moffat, J., et al. (2006). CellProfiler: image analysis software for identifying and quantifying cell phenotypes. Genome Biol 7, R100. 10.1186/gb-2006-7-10-r100.

53. Vahl, T.P., Ulrich-Lai, Y.M., Ostrander, M.M., Dolgas, C.M., Elfers, E.E., Seeley, R.J., D’Alessio, D.A., and Herman, J.P. (2005). Comparative analysis of ACTH and corticosterone sampling methods in rats. Am J Physiol Endocrinol Metab 289. 10.1152/AJPENDO.00122.2005.

