## Supplemental Figures for "Glucocorticoids desensitize hypothalamic CRH neurons to norepinephrine and somatic stress activation via rapid nitrosylation-dependent regulation of α1 adrenoreceptor trafficking"

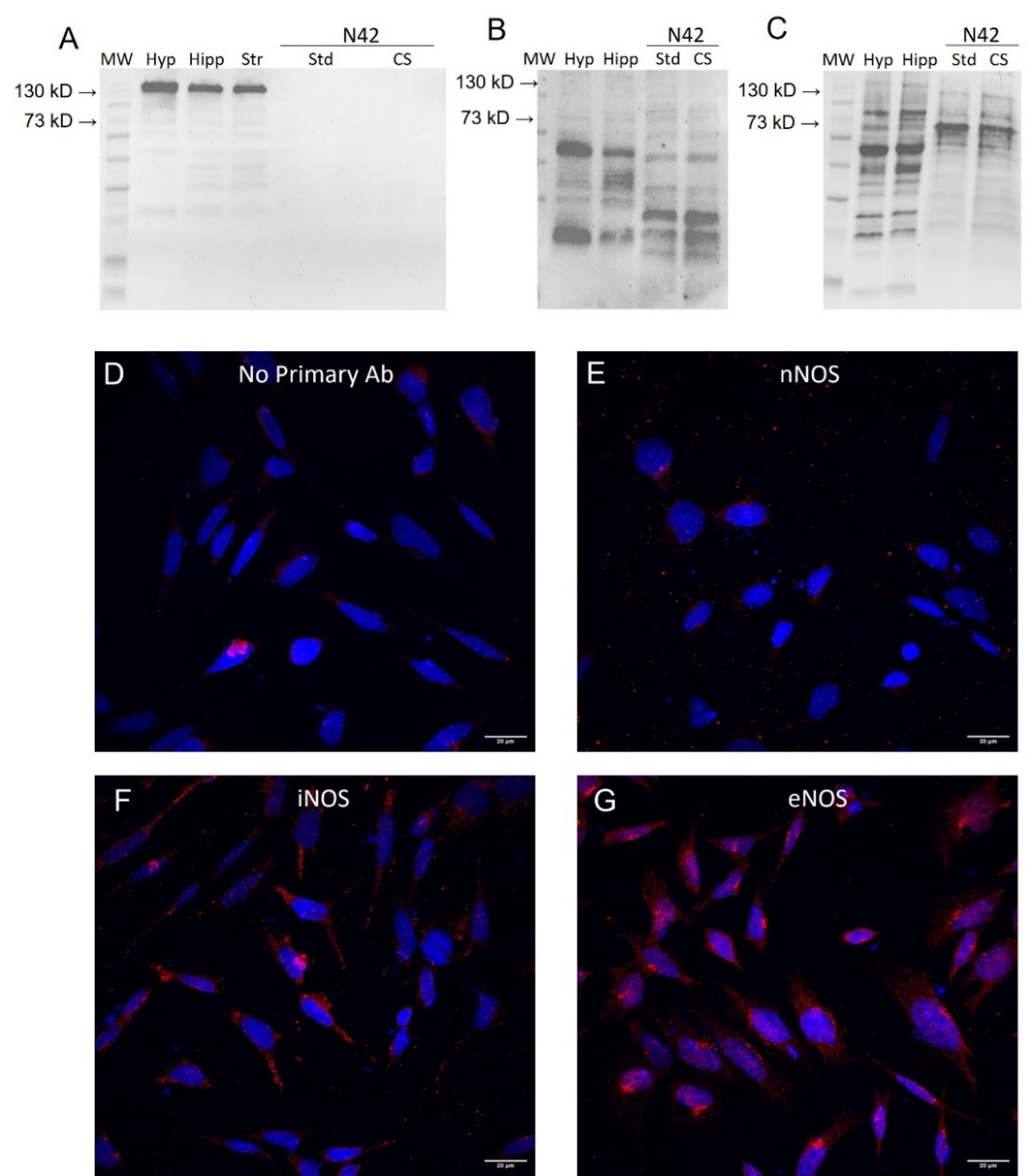
**Supplemental Figures**

**Supplemental Figure 1** (to go with Fig. 5). **NOS isoforms in N42 cells.** **A-C.** Western blots. **A.** nNOS (155 kD) was detected in brain hypothalamic (Hyp), hippocampal (Hipp), and striatal (Str) samples, but not in N42 cells cultured in either standard DMEM media (Std) or in DMEM with charcoal-stripped FBS (CS). **B.** iNOS (130 kD) was detected in hypothalamic (Hyp) and hippocampal (Hipp) tissue samples and in N42 cells. C. eNOS (140 kD) was detected in hypothalamic (Hyp) and hippocampal (Hipp) samples and in N42 cells. Std DMEM with 10% fetal bovine serum, CS DMEM with 10% charcoal-stripped fetal bovine serum. Brain samples: 20 µg/lane. N42 cells: 20 µg/lane (nNOS, iNOS), 30 µg/lane (eNOS). **D-G.** Immunocytochemistry of NOS isoforms in N42 cells. N42 cells were seeded in 24-well plates and cultured for ~36h. They were then switched to DMEM with charcoal-stripped fetal bovine serum and cultured for 16h to replicate assay conditions, then immunolabeled with (D) no primary antibody, (E) nNOS antibody, (F) iNOS antibody, and (G) eNos antibody. Blue = DAPI, Red = NOS immunolabel.

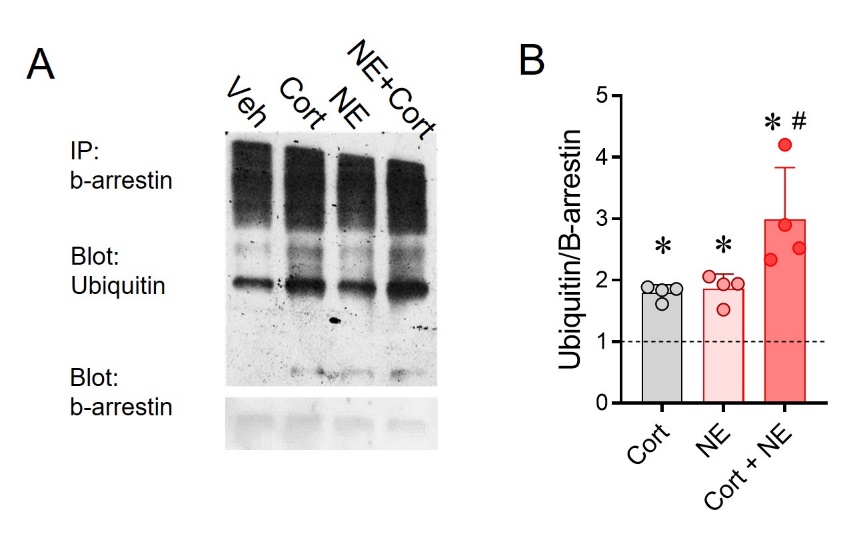

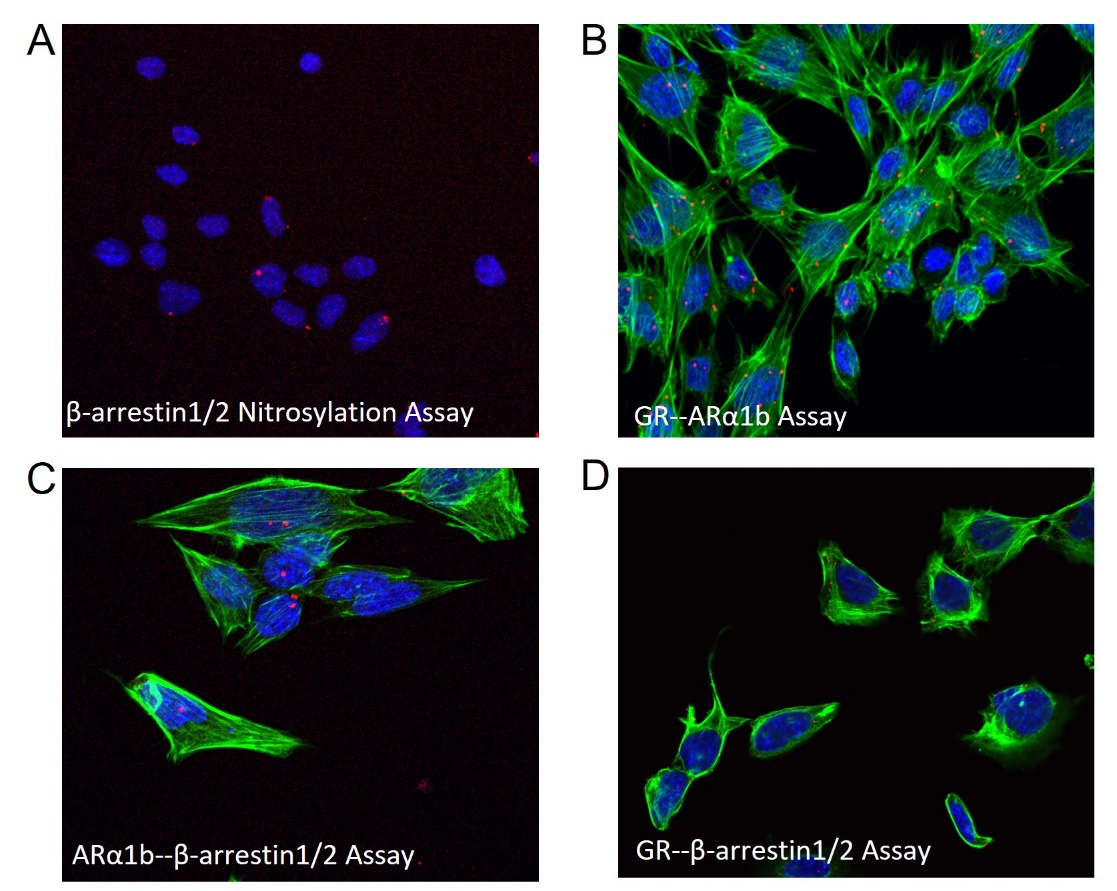

**Supplemental Figure 2. β-arrestin ubiquitination. A.** Immunoprecipitation of β-arrestin followed by blotting for ubiquitinated proteins. **B.** Quantification of ubiquitin signal normalized to blot for β-arrestin. 2 µM Cort caused a significant increase in ubiquitinated β-arrestin, comparable to 1 µM NE (ANOVA, * p<0.05, n=4). The effect of Cort and NE cotreatment is significantly more than NE alone (# p < 0.05 vs NE).

**Supplemental Figure 3.** **Proximity Ligation Assay single antibody controls.** Blue = DAPI staining of nuclei, red = PLA signal, green = phalloidin-FITC staining of F-actin. **A.** Β-arrestin1/2 antibody only. **B.** GR antibody only. **C.** Β-arrestin1/2 antibody only. **D.** GR antibody only.
