## Supplementary material for "Glucocorticoids desensitize hypothalamic CRH neurons to norepinephrine and somatic stress activation via rapid nitrosylation-dependent regulation of α1 adrenoreceptor trafficking": Table 1. Antibodies used

| **Antibody** | **Company** | **Dilution** |
| --- | --- | --- |
| Anti-Beta-arrestin 1/2 rabbit monoclonal | Cell Signaling Technology (#4672) | 1:100 |
| Anti-Biotin mouse monoclonal | Vector Labs (MB-9100) | 1:100 |
| Anti-GFP (B2) mouse monoclonal | Santa Cruz (sc-9996) | 1:100 |
| Anti-Glucocorticoid Receptor (M-20) rabbit polyclonal | Santa Cruz (sc-1004) | 1:300 |
| Anti-Glucorticoid Receptor (G-5) mouse monoclonal | Santa Cruz (sc-393232) | 1:50 |
| Goat anti-mouse IgG 594 | Abcam (ab96881) | 1:200 |
| Horse anti-mouse-HRP | Cell Signaling Technology (#7076) | 1:1000 |
| Anti-Myc mouse monoclonal | Sigma (M4439) | 1:200 |
| Anti-NOS1 (A-11) nNOS mouse monoclonal | Santa Cruz (sc-5302) | 1:200 (WB); 1:50 (ICC) |
| Anti-NOS2 (C-11) iNOS mouse monoclonal | Santa Cruz (sc-7271) | 1:200 (WB); 1:50 (ICC) |
| Anti-NOS3 (A-9) eNOS mouse monoclonal | Santa Cruz (sc-376751) | 1:200 (WB); 1:50 (ICC) |
